# CRISCross: Integration of Epigenetic Information for Deep-Learning-Based Prediction of CRISPR-Cas9 Off-Targets Across Cell Types

**DOI:** 10.64898/2026.10.09.757843

**Authors:** William Jobson Pargeter, Dominik Rabsch, Masako M. Kaufmann, Michael Uhl, Alexander Mitrofanov, Thomas Brox, Toni Cathomen, Van Dinh Tran, Rolf Backofen

## Abstract

CRISPR-Cas9 enables precise genome editing but remains limited by off-target effects, raising safety concerns and necessitating accurate prediction. Although epigenetic features influence Cas9 binding and cleavage, their predictive signals are often weak, highly variable, and context-dependent, limiting the effectiveness of existing machine learning approaches. Cross-cell-type settings are especially challenging: epigenetic landscapes differ substantially, and models trained on one cell type frequently fail to generalize. Here, we systematically investigate the role of epigenetic context in off-target prediction and demonstrate that its utility depends on modeling interactions with sequence features rather than treating it as an independent signal. Tabular and deep learning experiments demonstrate that epigenetic features provide a consistent but modestly complementary signal, and that late-stage fusion strategies are insufficient to effectively exploit this information. To address this, we introduce CRISCross, a cross-attention-based architecture that captures sequence-epigenetic interactions at the nucleotide level, alongside CRISPRAT, a synthetic pretraining strategy that learns transferable representations from genome-derived sgRNA-off-target pairs without relying on dataset-specific pretraining. CRISCross, incorporating AlphaGenome-derived chromatin accessibility signals, shows better predictive performance than sequence-only baselines across three independent test datasets spanning two cell types, demonstrating improved generalization across guide RNAs, datasets, and cellular contexts. This is particularly relevant for therapeutic applications involving cell types or patient-derived contexts that lack dedicated experimental datasets. In such settings, predicted chromatin accessibility may provide scalable contextual information when direct epigenomic profiling is unavailable or impractical. Together, these results show that accurate CRISPR off-target prediction benefits from interaction-aware integration of epigenetic context alongside generalizable pretraining strategies.

**Graphical Abstract:** 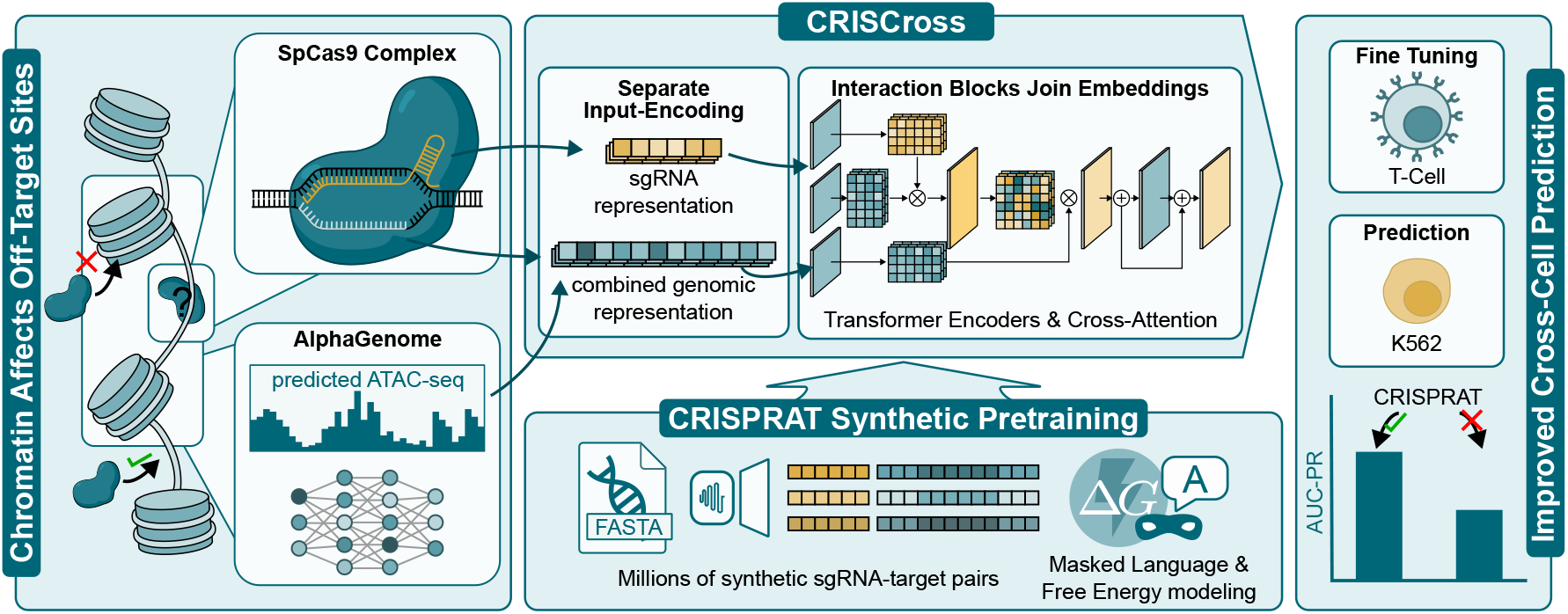

## Introduction

CRISPR-Cas9 has emerged as a widely used genome editing technology due to its simplicity, efficiency, and ease of programmability. Its applications span both basic research and clinical settings, including approved gene therapies for beta-hemoglobinopathies (1), ongoing clinical trials targeting inherited diseases (2, 3), and cancer therapies (4, 5). The CRISPR-Cas9 system consists of a single guide RNA (sgRNA) complexed with a Cas9 protein, which directs the nuclease to complementary genomic DNA sequences for targeted cleavage.

Despite its effectiveness, a key limitation restricting broader clinical application is the occurrence of off-target effects, including unintended sequence-level mutations and chromosomal rearrangements (6–9), caused by nuclease-mediated cleavage of genomic sites similar but not identical to the intended target. Experimental studies have shown that Cas9 can tolerate multiple mismatches between the sgRNA and DNA target sequence (8, 10). These additional edits raise safety concerns and highlight the importance of accurately predicting potential off-target sites, particularly in therapeutic contexts. Experimental assays have been developed to detect off-target activity, including cellular methods such as GUIDE-seq (10), DISCOVER-seq (11), and CAST-seq (7, 12), as well as in vitro techniques such as CIRCLE-seq (13) and CHANGEseq (14). Although these approaches provide high-sensitivity nomination of off-target sites, they are labor-intensive, costly, and difficult to scale across large numbers of sgRNAs and cell types. Furthermore, different assays often identify only partially overlapping sets of off-target sites, reflecting both biological variability and methodological differences. These limitations have motivated the development of computational approaches for off-target prediction.

Early in silico methods relied primarily on sequence alignment to identify genomic sites with a specified number of mismatches relative to the target sequence. Tools such as Cas-OFFinder (15) and SWOffinder (16) provide comprehensive lists of candidate sites but typically generate large numbers of false positives (17). Subsequent approaches incorporated heuristic rules (18) or thermodynamic models (19) to better approximate Cas9 binding and cleavage. More recently, machine learning and deep learning approaches have improved predictive performance by learning sequence features directly from data, including convolutional architectures (20–23) and transformer-based models with pretraining strategies (24–26). Most existing models focus primarily on the sequence pair formed by the sgRNA and the 23-nt candidate off-target region, which comprises the 20-nt guide-complementary sequence and the adjacent 3-nt PAM, without explicitly modeling the broader genomic context in which the site occurs (27). However, off-target activity is not determined solely by sequence similarity but is strongly influenced by local genomic context, including chromatin accessibility (28, 29) and nucleosome positioning (30). Importantly, these epigenetic signals are often weak, highly variable, and context-dependent, limiting their predictive utility when incorporated through naive integration strategies. This context dependence contributes to variability in off-target activity within a cell type and to even larger differences across cell types, where identical sgRNAs can exhibit divergent off-target profiles under different epigenetic conditions (31).

Experimentally determined epigenetic features provide a means to capture this contextual information. Chromatin accessibility can be measured using assays such as DNase-seq (32) and ATAC-seq (33), while histone modifications and transcription factor binding are typically profiled using ChIP-seq (34). However, experimentally generated epigenetic datasets are often noisy, assay-specific, and incomplete, limiting their direct applicability. Moreover, experimental profiling across diverse cell types is costly and time-intensive, restricting the availability of training data at scale. Recent sequence-based predictive models, including SEI (35), Enformer (36), and AlphaGenome (37), address these limitations by inferring celltype-specific epigenetic tracks directly from DNA sequence and a cell-type label, providing a scalable alternative with uniform genome-wide coverage.

Despite the potential of epigenetic context to improve offtarget prediction, it remains unclear how such signals should be effectively integrated into sequence-based models. This is particularly true in cross-cell-type settings where epigenetic landscapes differ substantially and models trained on one cell type frequently fail to generalize. In this work, we systematically compare experimental and predicted epigenetic features with respect to their ability to classify CRISPR off-target events, showing that although both are statistically informative, their predictive signals are weak and highly variable, limiting the effectiveness of naive integration strategies.

Based on these findings, we propose CRISCross, a framework for off-target prediction that explicitly models interactions between sequence and epigenetic context to capture cell-type-dependent activity. Unlike prior approaches that rely primarily on sequence features or incorporate epigenetic signals through simple concatenation, CRISCross employs a cross-attention mechanism that decouples sgRNA and target representations. This enables position-resolved interaction between sequence and epigenetic context at nucleotide resolution, rather than through summary statistics appended at the output layer. De-coupled encoding also removes the architectural constraint that the candidate site must match the sgRNA length, allowing genomic context beyond the 23-nt protospacer to be represented.

To address limitations of existing pretraining strategies, we further introduce CRISPRAT, a synthetic pretraining approach that learns transferable representations from largescale, genome-derived sgRNA–off-target pairs without relying on dataset-specific pretraining, with the aim of improving generalization to unseen sgRNAs, datasets, and cellular contexts. Through comprehensive evaluation, including crosscell-type prediction settings, we demonstrate that CRISCross achieves improved predictive performance compared to existing methods, highlighting the importance of interaction-aware modeling and transferable pretraining for accurate off-target prediction.

## Results and discussion

We present CRISCross, a CRISPR off-target prediction framework that separately encodes the sgRNA and genomic candidate sequence while integrating epigenetic context through cross-attention (Fig. 1; Methods). This design supports context-aware prediction beyond the fixed interaction window of paired encoding approaches. We further introduce CRISPRAT, a synthetic pretraining strategy designed to improve generalization across sgRNAs, datasets, and cell types.

**Fig. 1.**
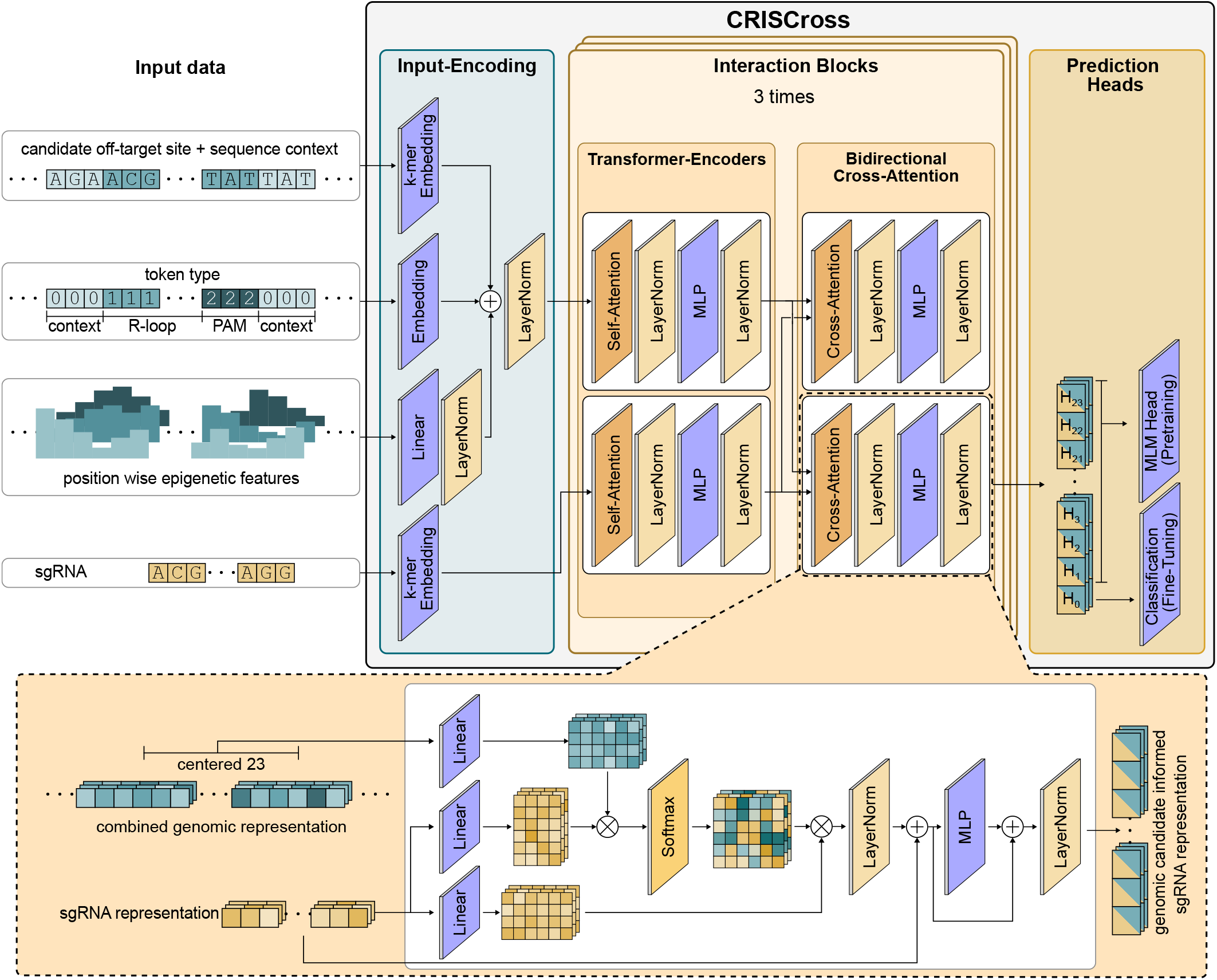
Overview of the CRISCross framework. Rather than treating the sgRNA and candidate off-target site as a single joint input (paired encoding), CRISCross encodes them as distinct sequences. Epigenetic signals are embedded alongside nucleotide features into a unified genomic representation, grounding predictions in both sequence composition and chromatin context. This design naturally accommodates off-target sites of varying length, freeing their representations from the constraints of the fixed sgRNA length. A series of interaction blocks takes sgRNA and genomic site representations as input, alternating between independent transformer encoding and cross-attention to facilitate bidirectional information exchange between the two representations and iteratively refine sequence interactions. The resulting sgRNA representation, enriched with contextual information from the genomic site, is used for downstream prediction. Task-specific MLP heads are used during training: a token-level head for the masked language modeling (MLM) objective during pretraining and a CLS-based classification head for off-target prediction during fine-tuning.

## CRISCross with contextual information and synthetic pretraining outperforms comparison models on AUCPR in cross-dataset, cross-cell-type prediction

We benchmarked CRISCross against existing CRISPR off-target prediction models by training on the T-cell dataset and evaluating on independent K562 and WTC-11 datasets. These test sets differ in cell type, sgRNA composition, candidate-site sampling, assay protocol, and class balance, making the evaluation both cross-cell-type and cross-dataset. Because the full model differs from the baselines in architecture, pre-training, and input features, we also report a sequence-only CRISCross variant under the same protocol, separating the AlphaGenome ATAC-seq contribution from that of the architecture and CRISPRAT pretraining.

CRISCross with AlphaGenome-predicted ATAC-seq achieved the highest AUC-PR on all three datasets: 0.587 on K562-I (from the DeepCRISPR paper (20)), 0.616 on K562-II (from the CRISPRoffT database (38)), and 0.545 on WTC-11, compared with 0.564, 0.563, and 0.466 for the best competing models, respectively (Fig. 2a). It also outperformed sequence-only CRISCross on all three datasets. Despite severe class imbalance, all models performed above the corresponding chance levels of 0.006, 0.004, and 0.003. On K562-II, the ATAC-seq variant outperformed sequence-only CRISCross for every guide and the other baselines for three of five guides (Fig. 2b). Corresponding per-guide results and across-seed standard deviations for K562-I and WTC-11 are provided in Suppl. Fig. S1. To quantify uncertainty arising from finite test-set sampling, we additionally calculated 95% stratified bootstrap confidence intervals for per-guide AUC-PR across all external guides (Suppl. Table S1). The confidence intervals generally became wider as the number of validated positive off-target sites decreased, highlighting increased sampling uncertainty for guides with fewer positives. For guides with very few positives, however, the bootstrap distributions can themselves be unstable or degenerate because only a small number of distinct positive observations are available for resampling; these estimates should therefore be interpreted cautiously. Precision at cutoff rank R was highest for CRISCross on K562-II and WTC-11 and second highest on K562-I, where CRISPR-OFFT performed best (Fig. 3a). Per-guide results and recall at 90% precision (Fig. 3b; Suppl. Table S1) show that no model dominates every setting, but CRISCross with ATAC-seq outperforms its sequence-only counterpart across almost all evaluated metrics and datasets.

**Fig. 2.**
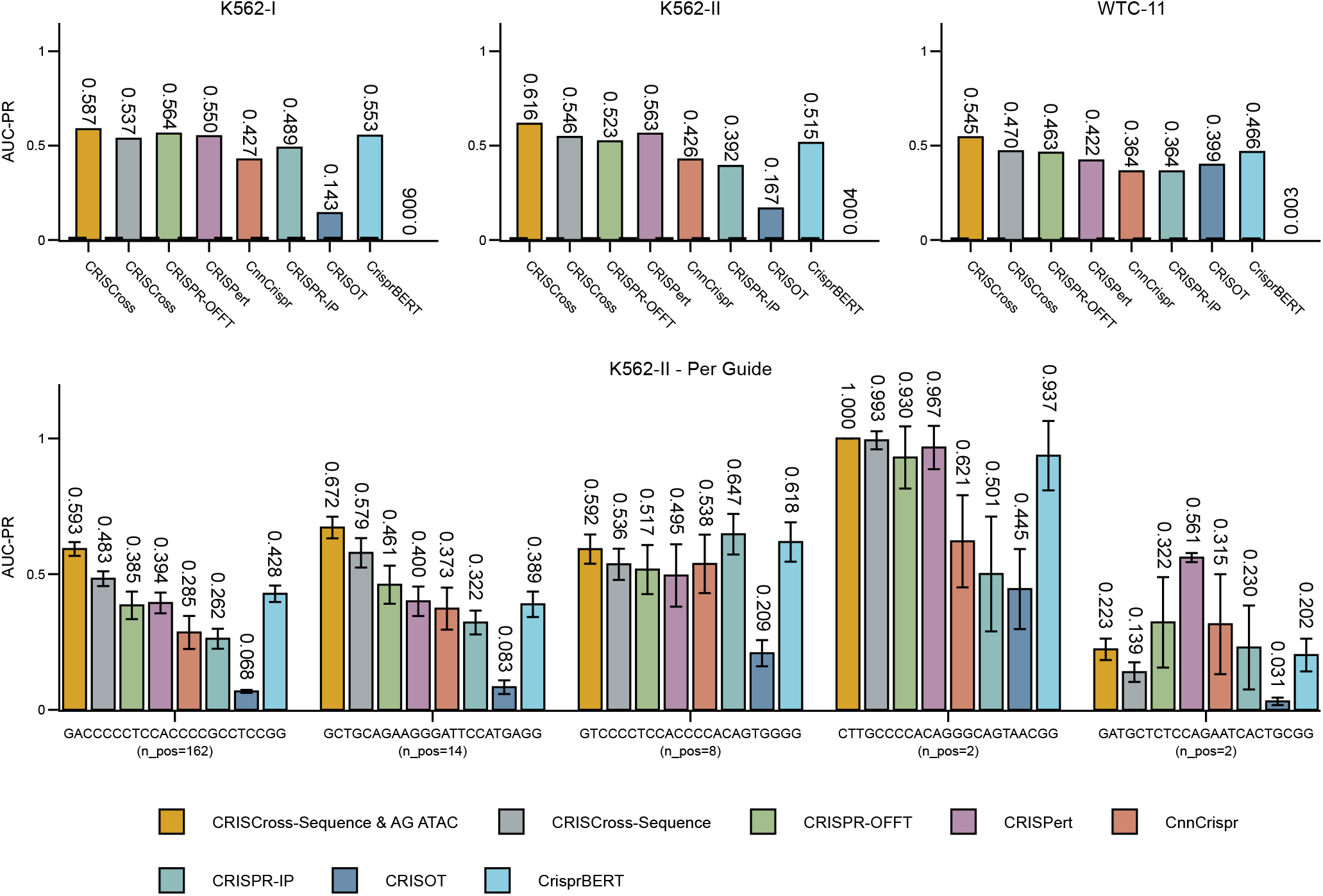
(a) Mean AUC-PR for each model on three held-out test sets (K562-I, K562-II, and WTC-11) averaged across all sgRNAs. The WTC-11 test set contains a single sgRNA. CRISCross is shown with both input configurations: sequence + AlphaGenome ATAC-seq (gold) and sequence only (gray). Dashed lines and adjacent values mark the AUC-PR expected by chance. (b) Per-sgRNA AUC-PR and across seed standard deviation on K562-II, with guides ordered by number of true-positive off-target sites (n_pos.).

**Fig. 3.**
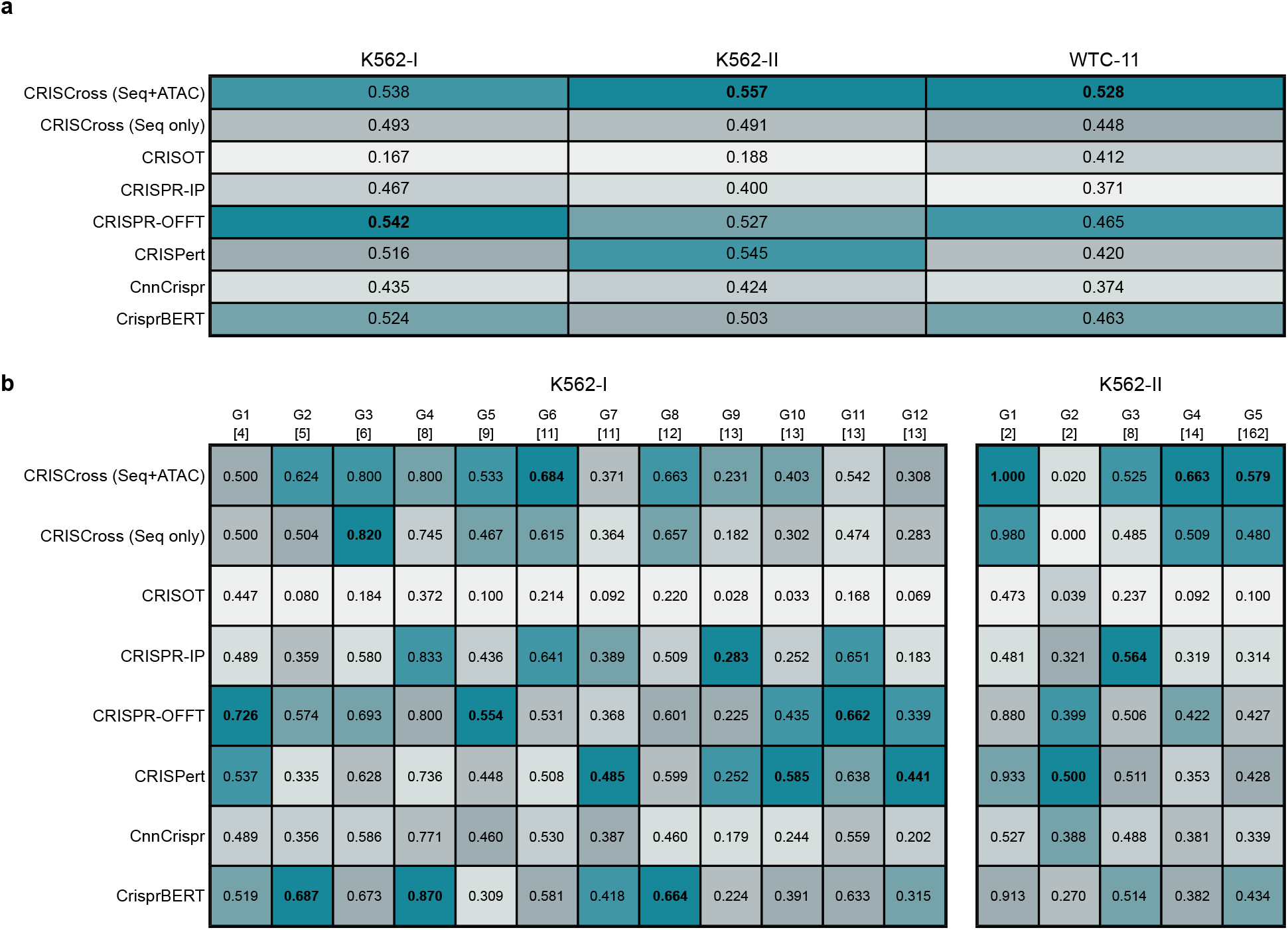
Precision at R across datasets and individual guides for off-target prediction models. (a) Mean precision at R - precision evaluated at each guide’s number of positives (R), averaged across all guides in a dataset - for eight models across three datasets: K562-I, K562-II, and WTC-11. CRISCross is shown with two feature configurations, sequence and AlphaGenome ATAC-seq (Seq+ATAC) and sequence only (Seq only). Bold values indicate the best-performing model for each dataset. Cell background colors indicate the relative ranking of models within each dataset (best to worst). (b) Precision at R for individual guides within each dataset. Guides are ordered by ascending R, with each guide’s R (number of positives) shown in brackets below its ID. A mapping from guide IDs to sequences is available in Suppl. Table S2. Bold values indicate the best-performing model for each guide. Cell background colors indicate the relative ranking of models for each guide.

## Sequence-context interaction modeling drives performance gains

Both the performance gains observed across the external test datasets and the final model architecture trace back to a series of prior analyses: evaluating epigenetic features in isolation, in tabular and deep learning models, and with explicit sequence–context interaction modeling.

### Epigenetic features provide informative but highly variable signals

To determine whether epigenetic features provide any predictive signal, we analyzed the distribution of epigenetic signals around off-target sites in an established GUIDE-seq T-cell dataset with matching epigenetic-marker tracks generated by high-throughput sequencing from (14). We evaluated their ability to separate positive and negative samples across different genomic context sizes, and further included AlphaGenome-predicted epigenetic features to compare their discriminative power with experimentally measured signals.

Although the mean distributions of epigenetic signals differ between positive and negative sites, they exhibit high variance (Fig. 4a). For most features, the mean lies outside the interquartile range, indicating pronounced skewness driven by outliers, and separability is further reduced when considering medians alongside interquartile ranges (Suppl. Fig. S2). This pattern is observed for both experimental and AlphaGenome-predicted features. While most features show statistically significant differences between classes according to the Mann-Whitney U test (Suppl. Table S3), the corresponding effect sizes, measured by the absolute rank-biserial correlation, remain small, indicating substantial overlap between positive and negative distributions.

**Fig. 4.**
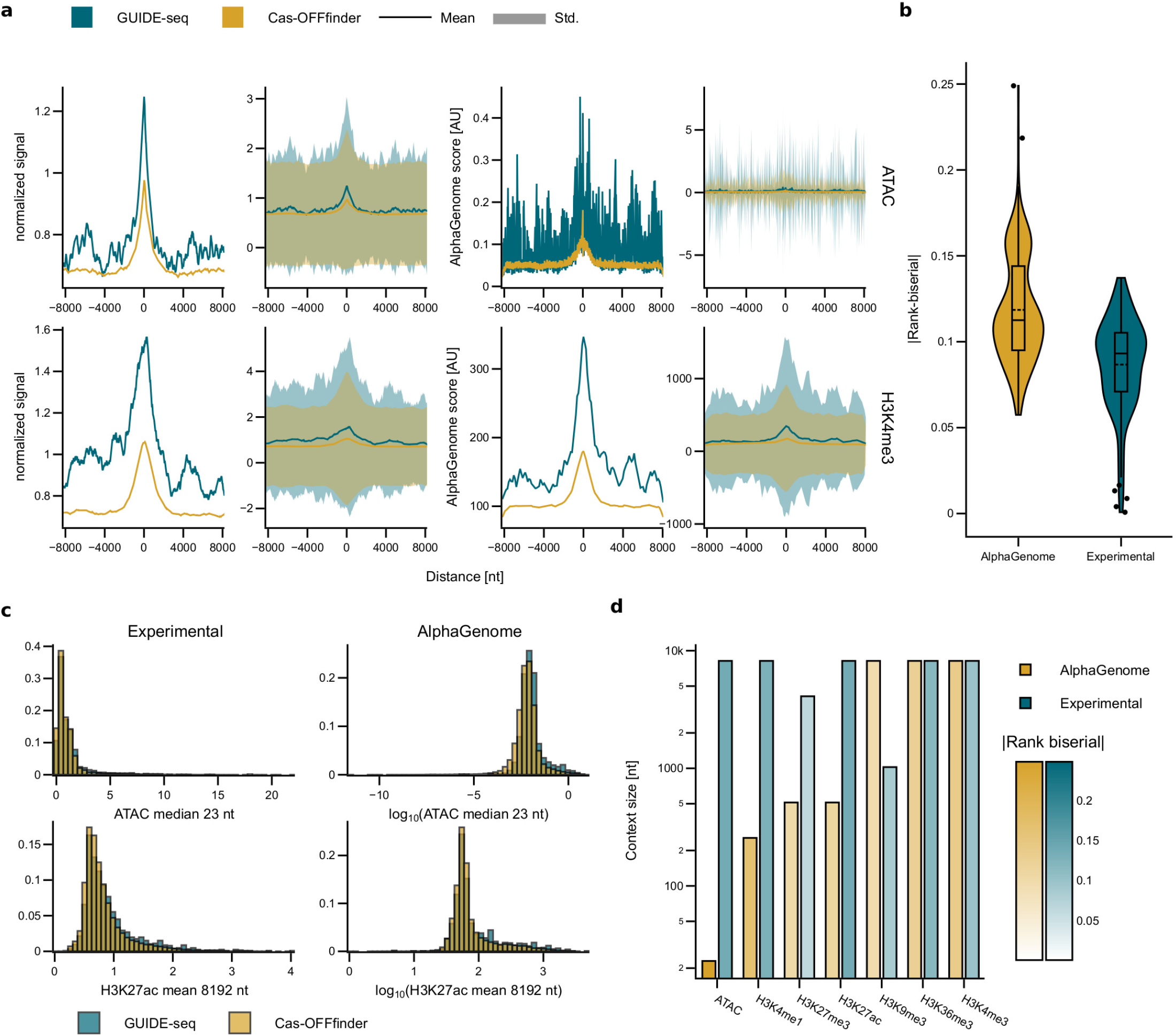
(a) Epigenetic feature values as a function of distance to the off-target site. Columns show mean and *±* one standard deviation ranges for both experimentally measured and AlphaGenome-predicted features. The substantial overlap of signals indicates limited separability between positive and negative sites. (b) Rank-biserial correlations for summary statistics (e.g. mean, minimum, maximum) derived from epigenetic features across different window sizes, separated into experimental and AlphaGenome feature sets. Only features with Holm–Bonferroni-adjusted *p <* 0.05 from two-sided Mann-Whitney U tests are shown. A value of 0 indicates no difference between groups, while positive values indicate higher feature values in GUIDE-seq-validated off-targets. (c) Distributions of top-ranked summary features from experimental data and AlphaGenome predictions. AlphaGenome ATAC-seq signal (median over 23 nt) shows the largest effect size among model-derived features, while experimental H3K27ac signal (mean over 8,192 nt) ranks highest among experimental features. GUIDE-seq (teal) and Cas-OFFinder (gold) distributions show highly significant differences, but moderate effect sizes lead to substantial overlap and limited visual separability. (d) Context size of the best-performing feature for each marker. The color scale represents the rank-biserial correlation value, where lighter colors indicate weaker discriminative power.

Several AlphaGenome-predicted features exhibit larger effect sizes than their experimental counterparts (Fig. 4b). The ATAC-seq median over a 23-nt window achieves the highest rank-biserial value of 0.249 (Fig. 4c), while the best-performing experimental feature, H3K27ac over an 8,192-nt window, ranks only 56th with a value of 0.137. Despite this difference in effect size, AlphaGenome-predicted accessibility still correlates reasonably well with experimentally measured chromatin accessibility (Pearson r around 0.5; Suppl. Fig. S3), indicating that the predicted tracks capture much of the same underlying biology rather than being an unrelated signal. Overall, effect sizes remain modest across both feature sets. We further examined the optimal context size for each epigenetic marker (Fig. 4d). For most experimental markers, the best-performing feature is observed at the maximum tested window size of 8,192 nt, whereas AlphaGenome-predicted features show more variation: ATAC-seq performs best at a 23-nt context and H3K4me1 at a 256-nt window. This difference is consistent with the underlying biology. ATAC-seq provides a direct, high-resolution readout of local chromatin accessibility, as Tn5 transposase preferentially integrates into nucleosome-free regions (39). Histone modifications, by contrast, reflect broader chromatin states and therefore provide a more indirect readout of accessibility (40). This likely contributes to their lower effect sizes and greater variability in optimal context size. The broader optimal windows observed for experimental features may additionally reflect their lower effective resolution compared to AlphaGenome-predicted signals, which capture more localized patterns.

The larger effect sizes for AlphaGenome-predicted features relative to experimental ones (Fig. 4b) likely reflect several compounding factors. Experimental tracks carry assay-specific noise, variable depth, and batch effects, whereas predicted tracks are smoothed, model-averaged estimates from large training compendia. Predicted tracks also provide uniform per-nucleotide coverage genome-wide, including in repetitive or low-mappability regions where experimental coverage is often sparse or unreliable.

Together, these results indicate that epigenetic features carry a statistically significant but weak and variable predictive signal. This raises the question of whether such signals can be effectively exploited by machine learning models, which we investigate in the following sections.

### Tabular models reveal dominant sequence effects with additional contributions from epigenetic features

To assess whether epigenetic features provide predictive benefit beyond sequence information, we evaluated TabPFN (41) in a T-cell leave-one-sgRNA-out (LOSO) setting. TabPFN is pretrained entirely on synthetic datasets rather than real data, and at inference the training split is supplied directly as context rather than used for gradient updates. Its pretrained weights therefore cannot have memorized any real off-target dataset. Because tabular models perform best with few features, we represented each sgRNA-off-target pair using the energy-based CRISPRoff score from (19) and a mismatch-position vector rather than one-hot or embedding encodings. The full feature set is described in the Supplementary Information.

Both sets of epigenetic features led to consistent improvements over the sequence-only baseline (Fig. 5a). The baseline model achieved an AUC-PR of 0.750, which increased to 0.763 with experimental features and 0.766 with AlphaGenome-derived features. These improvements are modest but statistically significant, and are consistent with our earlier feature-level analysis showing that AlphaGenomederived features tend to provide slightly stronger discriminatory signals.

**Fig. 5.**
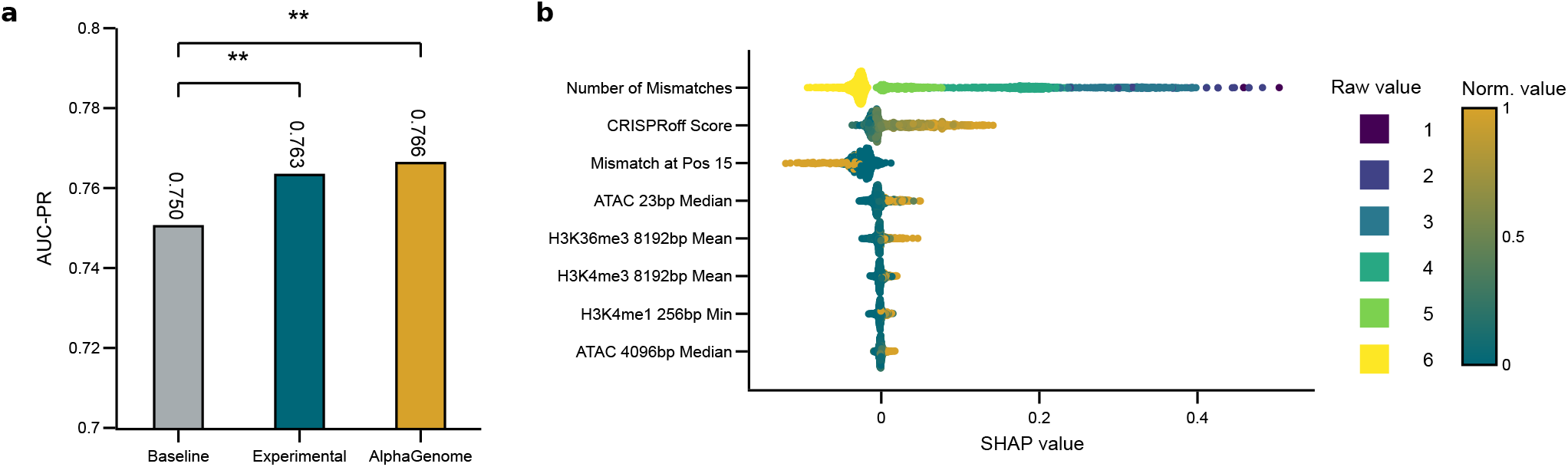
(a) Performance of TabPFN on the T-cell dataset using different feature sets. The baseline model includes no epigenetic features, while Experimental incorporates experimental epigenetic features and AlphaGenome incorporates AlphaGenome-derived features. Bars represent mean AUC-PR, error bars indicate standard deviation across splits, and values above bars denote mean performance. Asterisks indicate statistically significant improvements compared to the sequence-only baseline based on paired Wilcoxon signed-rank tests with Holm–Bonferroni correction (** *p <* 0.01). (b) SHAP analysis of the TabPFN model incorporating AlphaGenome-derived epigenetic features. The panel displays a subset of hand-selected features; the full SHAP results are available in the Supplementary Information. The number of mismatches exhibits a wide SHAP value distribution spanning both positive and negative values across samples, with a clear monotonic effect: lower mismatch counts consistently increased the predicted likelihood of true off-target activity. Similarly, at mismatch position 15 (a sensitive position; see text), the absence of a mismatch has little influence (SHAP values close to 0), whereas a mismatch reduces the prediction score for samples with this feature value. For CRISPRoff scores, corresponding to more favorable R-loop forming energies, low energies (normalized value close to 0) do not have much influence, but higher values are well correlated with prediction score. This implies that a higher R-loop score (equiv. to a lower free energy for the R-loop) is beneficial. Higher ATAC and H3K36me3 signals are likewise slightly beneficial; H3K36me3 is a hallmark of actively and recently transcribed genes.

To identify the features driving TabPFN predictions, we computed SHAP values (42) for the model incorporating AlphaGenome-derived features. Sequence-derived features dominated the top rankings (Fig. 5b). Mismatch count was most influential: sites with six mismatches received predominantly negative SHAP values, whereas strong positive contributions (SHAP *>* 0.2) were mainly associated with fewer than three mismatches. CRISPRoff score ranked second, reflecting more favorable guide-target binding energies and thus an increased predicted cleavage likelihood. Mismatch position 15 ranked third, though mismatches at nearby positions showed similar importance (Suppl. Fig. S4), with PAM-proximal positions being substantially more important than PAM-distal ones, consistent with the lower mismatch tolerance near the PAM (43). Epigenetic features contributed less: 23-nt ATAC-seq ranked 33rd of 57 features with a mean absolute SHAP value of 0.00684. Its directional effect was nonetheless consistent, with higher accessibility systematically shifting predictions toward true off-target classification, in line with the established role of chromatin accessibility in Cas9 binding (28, 29).

Together, these results show that sequence features dominate TabPFN predictions, while epigenetic features provide a weaker but consistent complementary signal. We next examine whether standard deep learning models can exploit this signal.

### Late merging of epigenetic features does not improve sequence-based models

To test whether standard deep learning models benefit from epigenetic information, we concatenated epigenetic summary statistics with the learned sequence representation at the final MLP layer of several existing off-target predictors, preserving their underlying sequence architectures (Fig. 6a). Across models, this late-merging strategy produced no statistically significant improvement (Fig. 6b). CRISPR-IP showed the largest nominal increase in AUC-PR, from 0.486 to 0.539, but gains were inconsistent and highly variable across random initializations and train-validation splits (Suppl. Fig. S5). These findings indicate that appending epigenetic features at the final prediction stage is insufficient and that more explicit sequence–context interaction modeling may be required.

**Fig. 6.**
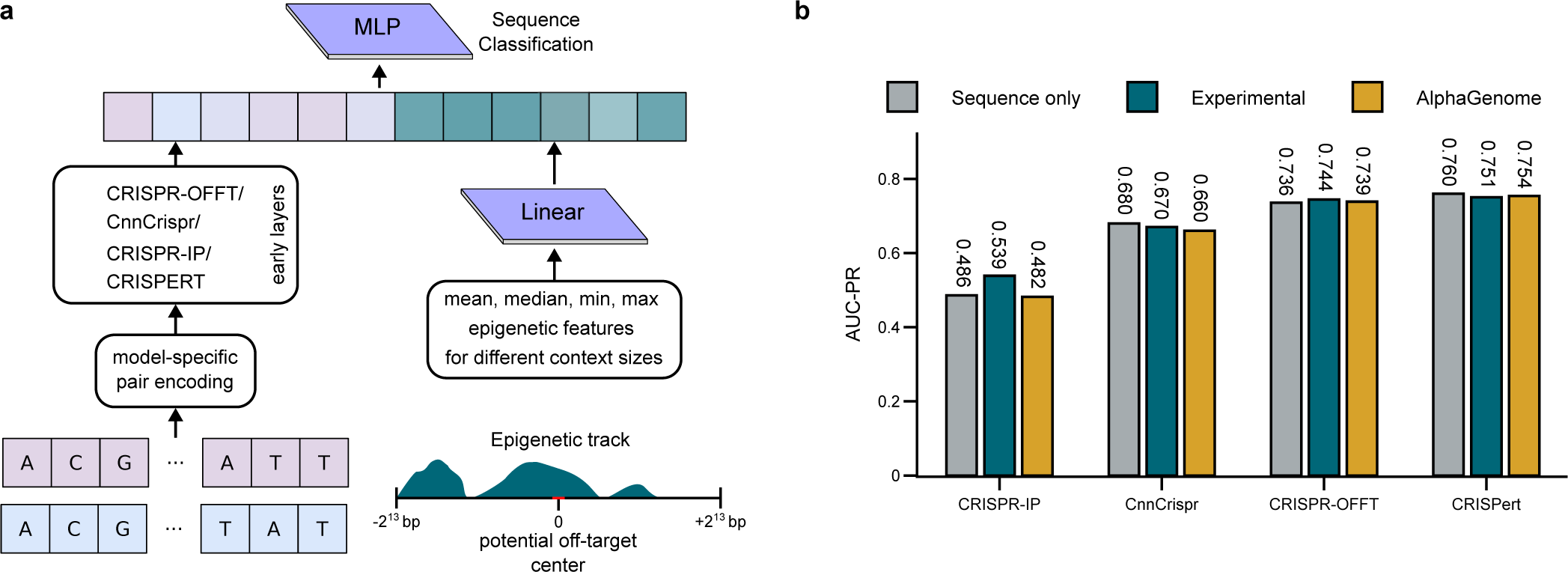
(a) Late-merging architecture, where sequence features are first processed to obtain latent representations, and summarized epigenetic features are concatenated at the final multilayer perceptron (MLP) layer. (b) Comparison of models trained using sequence features alone versus sequence features combined with epigenetic features through late merging. No statistically significant improvements were observed in Bonferroni-adjusted paired Wilcoxon signed-rank tests comparing models using experimental or AlphaGenome-derived features with the sequence-only model.

### A cross-attention architecture for integrating sequence with extended nucleotide-level epigenetic context

The previous results suggest that epigenetic features provide a complementary predictive signal, but that late-stage integration is insufficient to effectively leverage this information in standard deep learning models. We therefore hypothesized that epigenetic context must interact directly with sequence representations throughout the model to capture sequence-context relationships relevant to off-target activity.

To test this hypothesis, we evaluated a cross-attention-based architecture (CRISCross; see Methods) that separately encodes sgRNA and genomic context while enabling interaction between sequence representations and nucleotide-level epigenetic context (Fig. 1). Unlike joint encoding approaches used in most existing models, this design supports flexible input lengths and incorporation of extended genomic context beyond a fixed interaction window.

Because high-capacity architectures are difficult to train on small datasets, we additionally evaluated the effect of masked language model (MLM) pretraining. Among the evaluated baseline models, only CRISPert employs a pretraining strategy. We therefore compared CRISCross and CRISPert with and without pretraining under the same 23-nt context setting.

Without pretraining, both models performed below state-of-the-art levels (CRISCross: 0.031, CRISPert: 0.470). The substantially lower performance of CRISCross likely reflects the increased difficulty of learning sequence relationships without the inductive bias introduced by pairwise encoding. However, CRISCross exhibited a substantially larger improvement following MLM pretraining, reaching performance comparable to CRISPert (Fig. 7a). These results indicate that interaction-based architectures can effectively leverage pretraining while enabling integration of epigenetic context throughout the model.

**Fig. 7.**
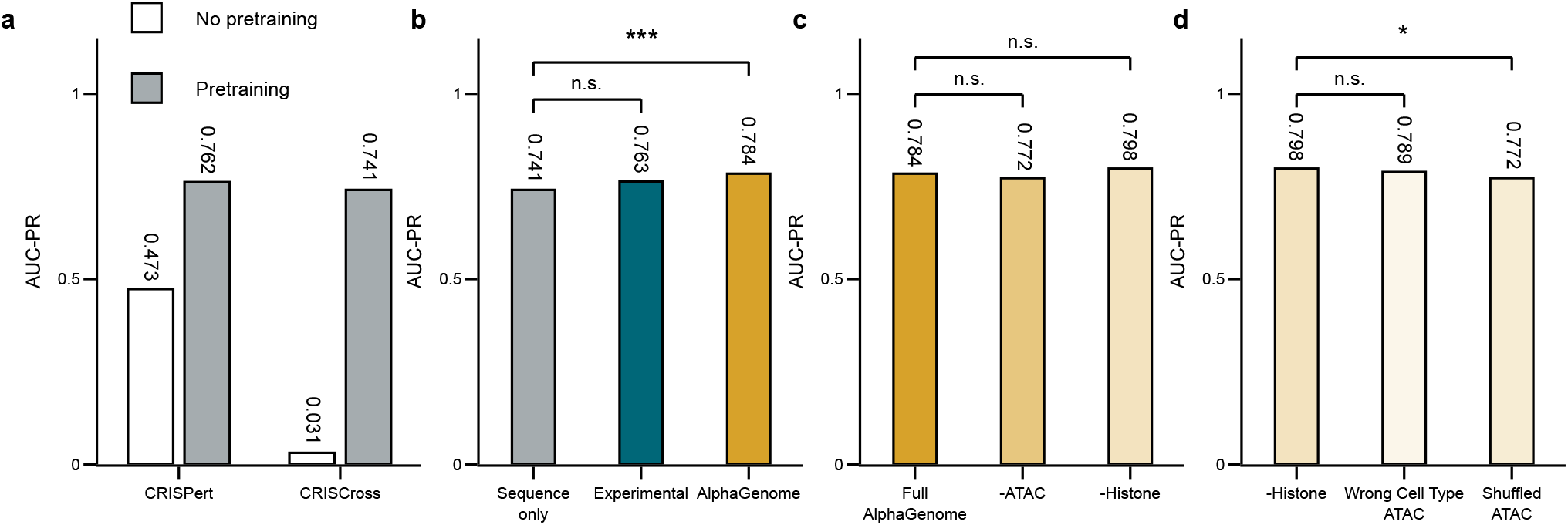
Performance comparison of CRISCross across feature sets and pretraining strategies. (a) Effect of masked language model (MLM) pretraining on AUC-PR for CRISPert and CRISCross, both evaluated at a 23-nt context window. (b) AUC-PR of CRISCross at 23-nt context size using sequence-only, experimental epigenomic (Experimental), or AlphaGenome-derived (AlphaGenome) input features. Each non-reference model is compared against the sequence-only baseline using a Wilcoxon signed-rank test with Holm-Bonferroni correction. (c) Ablation study of AlphaGenome input features at 23-nt context size. Models omitting ATAC-seq (-ATAC) or histone modification tracks (-Histone) are compared against the full AlphaGenome model. ATAC-seq features improve performance, while histone modification tracks are non-beneficial and their removal slightly improves results. (d) Impact of shuffled AlphaGenome ATAC input features at 23-nt context size. Models with fully shuffled tracks (Shuffled ATAC) or tracks from a mismatched cell type (Wrong Cell Type) are compared against the ATAC-only (-Histone) model. Fully shuffling tracks significantly degrades performance, whereas using tracks from the wrong cell type retains some predictive capacity. Significance levels: * *p <* 0.05, ** *p <* 0.01, *** *p <* 0.001, n.s. = not significant. P-values are Holm–Bonferroni-adjusted from paired Wilcoxon signed-rank tests across 17 sgRNA splits. Bar values indicate mean AUC-PR across cross-validation splits.

Together, these findings suggest that simple feature concatenation is insufficient for leveraging epigenetic information, and that effective integration may instead require explicit modeling of interactions between sequence and genomic context. Having established that sequence-only CRISCross performs on par with existing models when pretrained, we next evaluate whether incorporating epigenetic features into this interactionbased architecture yields measurable performance improvements.

### CRISCross effectively leverages predicted epigenetic context to improve performance

We incorporated epigenetic features into CRISCross and evaluated their impact by comparing models pretrained with either experimental or AlphaGenome-derived features against a sequence-only baseline, all at a context length of 23 nt.

The sequence-only model achieved an AUC-PR of 0.741. Incorporating experimental epigenetic features resulted in a modest and non-significant increase to 0.763, while AlphaGenome-predicted features led to a significant improvement, reaching an AUC-PR of 0.784 (Fig. 7b). These results indicate that epigenetic information provides a modest and source-dependent complementary signal. Publicly available experimental features did not significantly improve performance, whereas AlphaGenome-derived features produced a significant gain. We therefore do not conclude that measured epigenetic information is generally strongly predictive; rather, the benefit observed in this study is primarily associated with consistently represented, predicted epigenetic features, while sequence remains the dominant signal. These results are consistent with the feature-level analysis (Fig. 4b), which showed that predicted epigenetic features provide more consistent discriminatory signals in this dataset than experimental ones. Additional analyses evaluating larger context windows (128 and 512 nt) showed no significant improvement over the 23-nt baseline (Suppl. Results; Suppl. Fig. S6).

To assess the contribution of different epigenetic feature classes, we grouped AlphaGenome-derived features into histone modification markers, associated with broader chromatin states, and ATAC-seq features, which reflect local chromatin accessibility. Each ablated model was compared against the full AlphaGenome model (Fig. 7c). Although neither ablation resulted in a statistically significant difference, removing ATAC-seq features led to a noticeable decrease in performance, while excluding histone modification tracks resulted in a slight improvement. This suggests that within the 23-nt context window, ATAC-seq provides the most informative signal, while histone markers contribute limited additional information. This is consistent with the earlier feature analysis showing that histone signals are most discriminative at substantially larger genomic windows (Fig. 4d). Based on these ablation results, the final cross-cell-type CRISCross model uses only AlphaGenome-derived ATAC-seq features. Histone modification tracks were excluded because they did not improve performance.

To test whether CRISCross learns biologically meaningful, site- and position-specific epigenetic signal rather than simply benefiting from an extra numeric input channel, we evaluated two negative controls at 23-nt context size: AlphaGenome ATAC-seq tracks fully shuffled across sites and positions in the T-cell dataset, and ATAC-seq tracks predicted for an incorrect cell type (K562) (Fig. 7d). Both reduced performance relative to the intact, correct-cell-type model (0.798), with full shuffling causing a larger, significant drop (to 0.772) and the wrong cell type a smaller, non-significant one (to 0.789). To further characterize these controls, we compared the mean ATAC-seq signal across each 23-nt window. The T-cell and K562 tracks were strongly correlated across the 51,906 candidate sites (Pearson *r* = 0.809, *p <* 10^*−*300^), whereas the shuffled T-cell signal showed essentially no correlation with the original T-cell signal (*r* = −0.004, *p* = 0.366). The strong similarity between the T-cell and K562 accessibility profiles may partly explain the relatively small performance difference between the matched and mismatched cell-type conditions. Full shuffling still outperformed the sequence-only baseline (0.741, Fig. 7b). This difference is likely due to our shuffling approach mirroring a noise-injection technique that is known to improve model performance (44). Together, these results indicate that CRISCross benefits from preserving structured site- and position-specific chromatin accessibility and suggest an additional advantage of using cell-type-matched chromatin information, although this latter effect did not reach statistical significance.

Overall, these results demonstrate that CRISCross effectively leverages predicted chromatin accessibility signals when epigenetic features are integrated throughout the sequence representation learning process. Having shown that the model benefits from epigenetic context within a single cell type, we next examine whether this advantage extends to generalization across datasets and cell types.

### CRISPRAT pretraining improves generalization to unseen sgRNAs and datasets

While pretraining is critical for high-capacity models on small datasets, existing approaches such as CRISPert rely on dataset-specific pretraining, which limits generalization to unseen sgRNAs and cell types. To address this, we developed CRISPRAT (CRISPR Artificial preTraining; see Methods), a synthetic pretraining strategy that samples genomic loci directly from the human reference genome and generates artificial sgRNA sequences by introducing up to six mismatches, approximating the mismatch distribution observed in fine-tuning datasets (Fig. 8a).

**Fig. 8.**
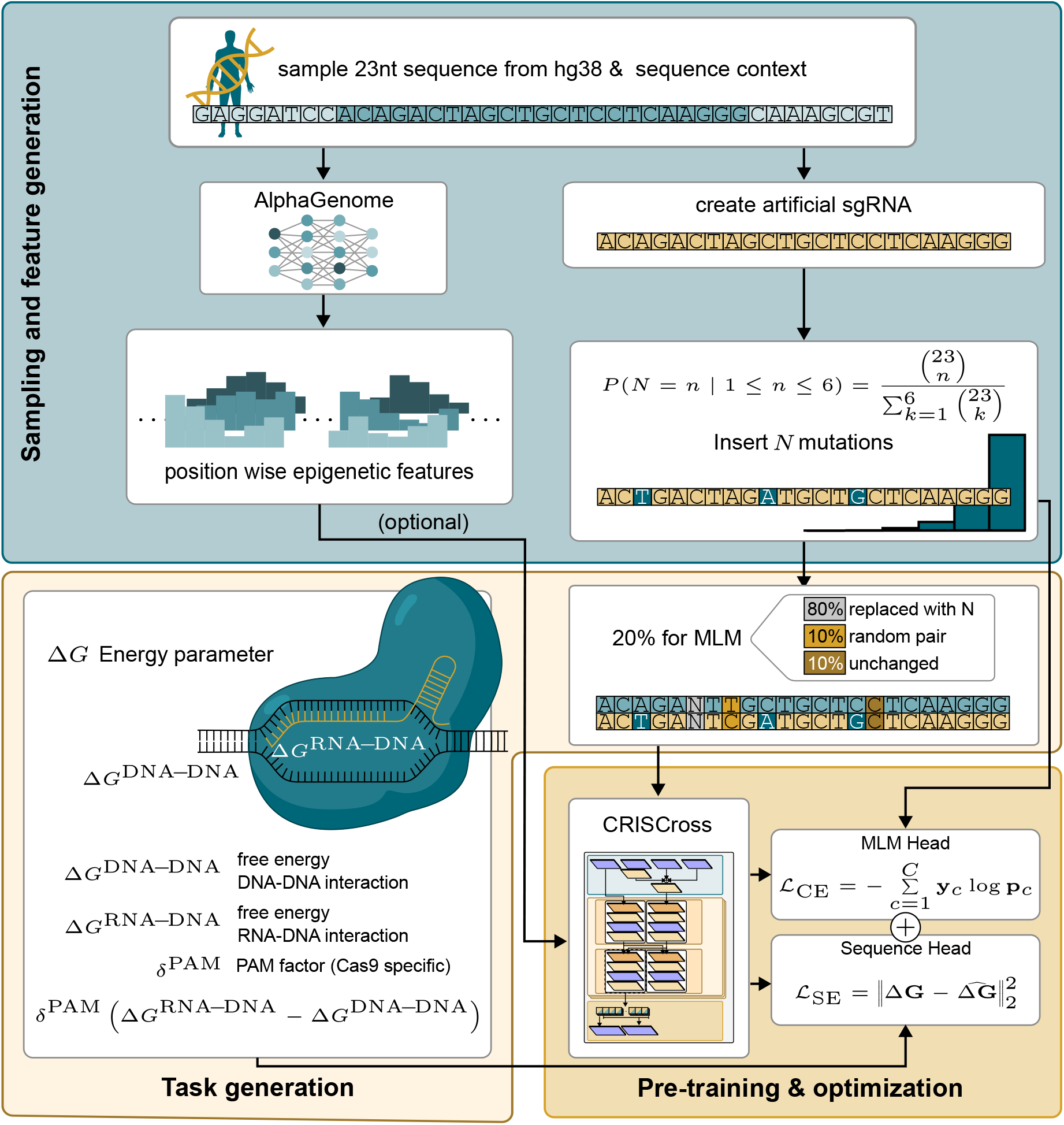
Overview of the CRISPRAT pretraining strategy. CRISPRAT trains CRISCross by generating artificial sgRNAs from genomic target sequences with controlled mismatch distributions and using them for masked language modeling and thermodynamic regression. Optionally, the model uses position-wise epigenetic features predicted by AlphaGenome.

We assessed the generalization ability of CRISPRAT-pretrained CRISCross models across within-dataset and cross-dataset settings (Fig. 9). In the T-cell LOSO evaluation, no significant difference was observed between CRISPRAT and dataset-specific pretraining, indicating that synthetic pre-training matches the performance of task-specific pretraining within the same dataset. In contrast, when evaluated on the unseen K562-I dataset, CRISPRAT pretraining resulted in a substantial improvement, achieving an AUC-PR of 0.783 compared to 0.569 for the model pretrained on the T-cell dataset. These results demonstrate that CRISPRAT improves robustness to dataset shifts without requiring access to target dataset sequences during pretraining.

**Fig. 9.**
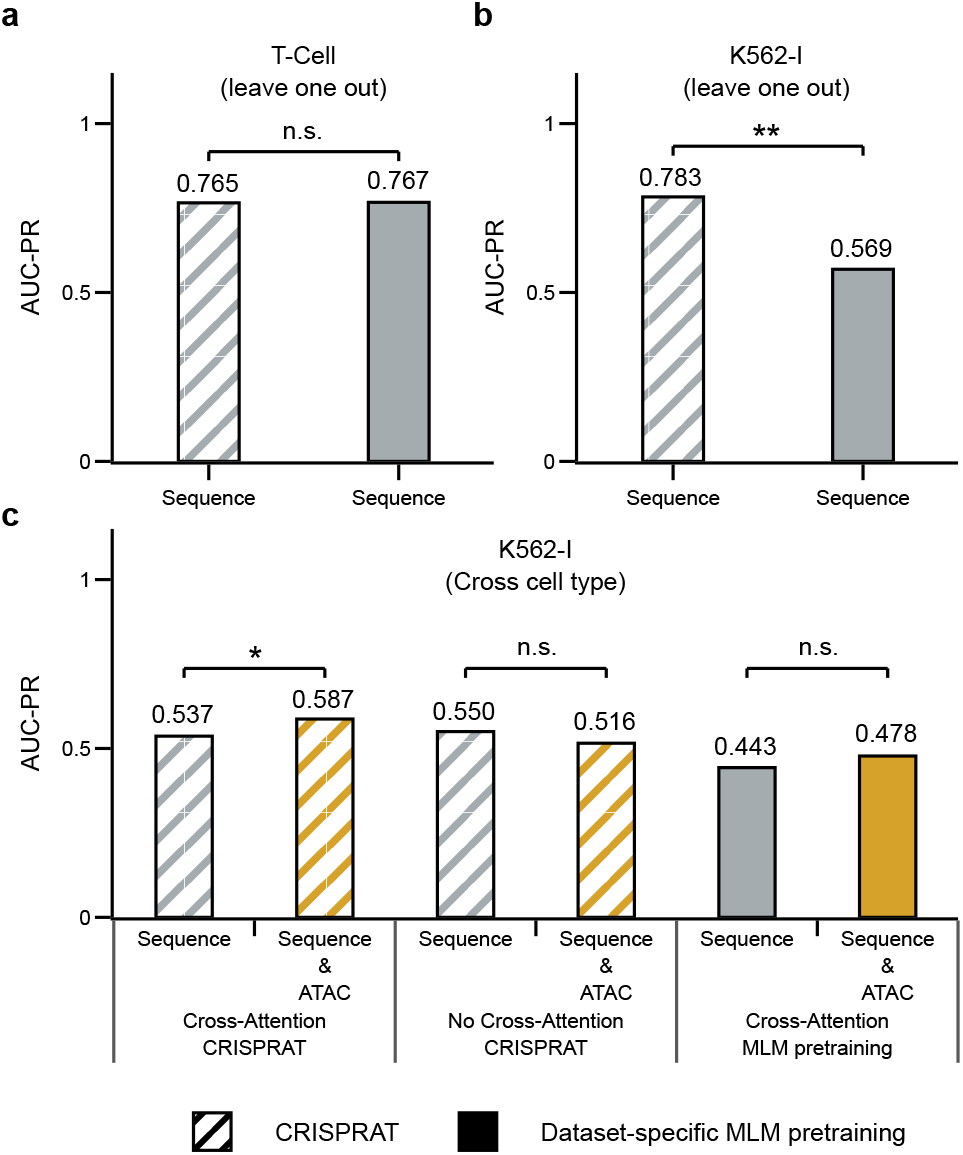
(a) It can bes observed that when models are fine-tuned on datasets containing sgRNAs that were already present during pretraining, there is no significant difference between the pretraining strategies. (b) In contrast, when the models are fine-tuned on datasets containing previously unseen sgRNAs, CRISPRAT significantly outperforms traditional dataset-specific MLM pretraining. (c) In the lower panel for cross-cell-type evaluation, cross-attention and CRISPRAT are required to effectively use sequence and ATAC signals. The lower AUC-PR values in the cross-dataset, cross-cell-type evaluation are expected due to the increased difficulty of transferring across distinct cell types and experimental conditions. Statistical significance is indicated by asterisks (* *p <* 0.05, ** *p <* 0.01). Significance was assessed using paired Wilcoxon signed-rank tests across sgRNAs, and all p-values were adjusted using the Holm-Bonferroni method.

We next evaluated whether incorporating epigenetic information during pretraining further improves cross-dataset and cell-type generalization. To this end, we compared CRISPRAT models pretrained using sequence alone against models additionally incorporating AlphaGenome-derived ATAC-seq signals, with both subsequently fine-tuned on the T-cell dataset and evaluated on K562-I.

The model incorporating ATAC-seq signals during pretraining achieved a substantially higher AUC-PR of 0.587 compared to 0.537 for the sequence-only model, indicating that integrating epigenetic context during pretraining helps the model learn regulatory representations that transfer more effectively to the independent K562-I dataset, which differs from the training data in cell type as well as sgRNA composition, assay protocol, and class balance. This finding was robust to alternative negative-sampling strategies during fine-tuning, including repeated random subsampling and hard-negative sampling (Suppl. Results).

We then investigated to what extent CRISPRAT contributed to CRISCross’s performance in a cross dataset, cross-cell-type setting on the K562-I dataset. To this end, we compared the full model with two ablations: (i) CRISCross pretrained with dataset-specific MLM instead of CRISPRAT, and (ii) CRISCross with cross-attention replaced by simple additive fusion of the two representations, each evaluated with and without AlphaGenome-derived ATAC-seq features (Fig. 9). Interestingly, replacing CRISPRAT pretraining with dataset-specific MLM reduced performance (AUC-PR 0.443 Sequence vs. 0.478 Sequence+ATAC, not significant), as did substituting cross-attention with additive fusion (0.550 vs. 0.516, not significant). This suggests that the performance gain from AlphaGenome ATAC-seq features depends specifically on the combination of CRISPRAT pretraining and cross-attention fusion, rather than either component alone.

Taken together, these results support CRISPRAT as a synthetic genome-based pretraining framework that improves generalization across sgRNAs, datasets, and cell types. More broadly, they demonstrate that epigenetic context contributes not only to within-dataset prediction but also to generalization across datasets and cellular contexts when incorporated during pretraining.

## Conclusion

In this work, we investigated the role of epigenetic context in CRISPR-Cas9 off-target prediction and demonstrated that, although epigenetic features are statistically informative, their predictive signal is weak, highly variable, and difficult to exploit through naive integration strategies. Systematic evaluation showed that late-stage fusion of epigenetic features into existing sequence-based models consistently fails to yield significant improvements, motivating a more principled approach to sequence-context integration.

To address this, we introduced CRISCross, a cross-attention-based architecture that models interactions between sequence and epigenetic representations at the nucleotide level, enabling effective integration of contextual signals throughout the model. Alongside this, we proposed CRISPRAT, a synthetic pretraining strategy based on genome-derived sgRNA-off-target pairs that improves generalization across sgRNAs, datasets, and cell types without requiring dataset-specific pre-training.

Through comprehensive evaluation across three independent test datasets spanning two cell types, CRISCross incorporating AlphaGenome-derived chromatin accessibility signals showed improved predictive performance over sequence-only baselines overall. These results demonstrate that predicted epigenetic features can provide consistent and transferable improvements over sequence-only approaches when integrated through cross-attention-based interaction modeling.

Practically, CRISCross could help prioritize candidate offtarget sites in cell types lacking dedicated experimental data, reducing reliance on costly assays like GUIDE-seq for initial screening. However, cross-cell-type performance remains well below within-cell-type levels and was evaluated on limited dataset pairs, so this should be viewed as a promising direction rather than a validated substitute for experimental profiling.

More broadly, in our evaluation the AlphaGenome tracks were the more useful model inputs, which suggests sequence-to-epigenome predictions may be a practical source of regulatory context for off-target prediction even where experimental data exist. This reflects their utility as features in this setting rather than greater biological accuracy than the measured tracks.

Together, these findings indicate that accurate CRISPR off-target prediction benefits not only from incorporating epigenetic context but also from learning its interaction with sequence features through interaction-aware architectures and transferable pretraining strategies. More broadly, the proposed framework provides a foundation for integrating regulatory context into CRISPR off-target prediction and improving robustness across diverse cellular environments.

## Methods

We describe the datasets, evaluation protocols, and computational methods used to develop and assess CRISCross and CRISPRAT.

## Datasets and evaluation protocol

### Datasets

Four experimentally validated CRISPR off-target datasets were used in this study to evaluate model generalization within and across cell types.

#### The T-cell dataset

Derived from GUIDE-seq data reported by Lazzarotto et al. (14), containing off-target sites for 78 sgRNAs measured in human primary CD4^+^/CD8^+^ T cells. After filtering low read-count sites and sgRNAs with fewer than 10 off-targets, 17 sgRNAs with 1,266 positive off-target sites were retained. Negative samples were generated using Cas-OFFinder by scanning for all sites with *≤* 6 mismatches, followed by random sampling to achieve a 1:40 class imbalance, resulting in 50,604 negative samples. To assess robustness to negative sampling, we generated six independent random negative datasets, plus one hard-negative dataset using the same positive sites. For each positive site, hard negatives were the 40 closest non-target sites based on mismatch count and position, rather than random selection.

#### The K562-I dataset

Derived from the DeepCRISPR study (20), comprising 12 sgRNAs with 118 experimentally validated off-target sites as positives and 18,303 candidate sites as negatives.

#### The WTC-11 dataset

Derived from the CRISPRoffT database (38) with one sgRNA from WTC-11. Once again, negative samples were generated using Cas-OFFinder by scanning for all sites with *≤* 6 mismatches, without subsampling.

#### The K562-II dataset

Derived from the CRISPRoffT database (38) with five sgRNAs from GUIDE-seq/DISCOVER-seq. Negative samples were generated similarly to those in the WTC-11 dataset.

One sgRNA from the K562-II dataset overlaps with the T-cell dataset, although the identified positive off-target sites differ. No sgRNAs from either of the other two datasets overlap with the T-cell dataset.

### Feature extraction and preprocessing

#### Epigenetic features

Experimental epigenetic features, including chromatin accessibility (ATAC-seq) and histone modification tracks (H3K4me3, H3K27me3, H3K4me1, H3K9ac, H3K27ac, H3K36me3), were obtained as big-Wig files from the Gene Expression Omnibus (accession GSE149363). Predicted epigenetic features were generated using AlphaGenome, which infers cell-type-specific epigenetic tracks directly from DNA sequence and a cell type label. Predictions were generated with a window size of 2^14^ bp using ontology terms CL:0000084 (T-cell), EFO:0002067 (K562), and EFO:0009747 (WTC-11). Detailed preprocessing procedures for both feature types are provided in the Supplementary Methods.

#### Energy features

Energy-based features for DNA-DNA, RNA-DNA, and RNA-RNA interactions were computed as described in (19). While these features were used for tabular models, RNA-RNA interaction energies were excluded from CRISPRAT to reduce computational cost.

### Evaluation setup

#### Evaluation protocol

We evaluated model performance under two settings. In the leave-one-sgRNA-out (LOSO) setting, all off-target sites associated with one sgRNA were held out as a test set while the remaining data were used for training; this process was repeated for all sgRNAs and mean performance across splits was reported. In the cross-cell-type setting, models were trained on the T-cell dataset and evaluated on the K562-I, K562-II, and WTC-11 datasets as independent test sets. For each test dataset, performance was computed for each sgRNA and then averaged across all sgRNAs in that dataset. All experiments were repeated across multiple random seeds, and results are reported as averages across runs, using 10 seeds for LOSO and 25 for cross-cell-type evaluation. **Evaluation metrics:** We report the area under the precision-recall curve (AUC-PR) as the primary evaluation metric. We additionally report precision at R, where R is the number of positives per sgRNA. Statistical significance of performance differences was assessed using the paired Wilcoxon signed-rank test with Holm-Bonferroni correction, treating individual guide RNAs as paired observations. For feature-level analysis, we used the Mann-Whitney U test and rank-biserial correlation to quantify effect sizes.

#### Feature attribution

We used SHAP values (42) to assess feature contributions to model predictions. A detailed description is provided in the Supplementary Methods.

### CRISCross framework

CRISCross is a neural architecture designed to model interactions between sgRNA sequences and genomic candidate sites by integrating nucleotide sequence, epigenetic signals, and positional region annotations. It consists of three main building blocks: input encoding, interaction blocks, and prediction heads (Fig. 1).

#### Input encoding

The sgRNA and genomic candidate sequences are tokenized independently into overlapping trinucleotides (3-mers) and mapped to embedding vectors through a learned token embedding layer. A special classification (CLS) token is prepended to the sgRNA sequence, whose final representation is used as a global summary for downstream prediction.

Epigenetic context is incorporated exclusively into the genomic candidate representation. Nucleotide-wise epigenetic feature values are projected into the same embedding space using a linear layer. Each position in the genomic candidate sequence is additionally assigned a region label encoding its functional role: PAM positions are labeled 1, sgRNA binding site positions are labeled 2, and all other positions are labeled 0. Region labels are mapped to learned embedding vectors. The epigenetic and region embeddings are combined with the genomic sequence representation via element-wise summation, yielding a unified genomic-epigenetic-region representation.

#### Interaction blocks

The core of CRISCross consists of a stack of *N* = 3 interaction blocks, each integrating independent sequence encoding with cross-sequence interaction. Within each block, the sgRNA and genomic-epigenetic representations are first processed separately using dedicated Transformer encoder layers. The resulting representations are then coupled through bidirectional cross-attention restricted to the central 23-nucleotide region of the genomic candidate sequence, corresponding to the sgRNA binding site. This produces updated representations for both the sgRNA and the central genomic-epigenetic region, which are concatenated with the unchanged flanking representations to restore the original sequence length. Both representations are propagated through subsequent blocks, with the final sgRNA CLS representation used for downstream prediction.

#### Prediction heads

Two MLP prediction heads are used depending on the training objective: a token-level MLM head applied to sgRNA representations during pretraining to reconstruct masked tokens, and a classification head applied to the CLS representation during fine-tuning to output a scalar off-target likelihood.

### Synthetic pretraining (CRISPRAT)

CRISPRAT (CRISPR Artificial preTraining) is a synthetic pretraining strategy designed to learn generalizable representations for CRISPR off-target prediction without relying on dataset-specific pretraining. Rather than exhaustively enumerating candidate off-target sites using alignment-based search tools, genomic loci are sampled directly from the hg38 reference genome and artificial sgRNA sequences are generated by introducing sampled mismatches into the target sequence (Fig. 8a). To approximate the mismatch distribution observed in fine-tuning datasets, the number of mismatches *N* is sampled according to a combinatorial distribution:

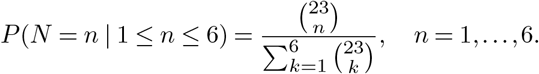

CRISCross is pretrained using a masked language modeling (MLM) objective in which 20% of nucleotide tokens within the interaction window are randomly masked and the model is trained to reconstruct the original sequence. An auxiliary regression objective is additionally used to predict an energy-based parameter reflecting the thermodynamic properties of R-loop formation (19), introducing CRISPR-specific inductive bias into the pretraining process. When epigenetic features are available, AlphaGenome-derived ATAC-seq signals are incorporated into the genomic candidate representation during pretraining, enabling the model to learn sequence-epigenetic interactions across large-scale synthetic genomic contexts.

### Baselines and implementation details

#### Baseline models

We compared CRISCross against six established CRISPR off-target prediction models representing diverse architectural designs: CnnCrispr (22), CRISPR-OFFT (21), CRISPR-IP (23), CrisprBERT (45), CRISOT (46), and CRISPert (24). CnnCrispr, CRISPR-OFFT, CrisprBERT, CRISOT, and CRISPR-IP were implemented according to their published configurations. For CRISPert, a reduced version with six instead of twelve stacked transformer blocks was used to better match the scale of the available training data. Preliminary experiments showed comparable performance with six and twelve transformer blocks (Suppl. Fig. S7). We therefore used the six-block configuration to reduce unnecessary model capacity for the short input sequences and the limited dataset size. All models were trained and evaluated using the same data splits and evaluation protocols described above. For experiments incorporating epigenetic features, late-merging architectures were extended by concatenating summary statistics of epigenetic signals at the final MLP layer using identical feature sets and preprocessing procedures.

#### Implementation details

CRISCross was implemented in PyTorch. Key hyperparameters and training configurations, including learning rate schedules, batch sizes, class imbalance handling, and model configurations, are provided in the Supplementary Methods.

#### Limitations

Several factors limit the scope of CRISCross. First, the model was trained and evaluated exclusively on SpCas9 in human cells, so it remains unclear how well the model or epigenetic features generalize to other Cas nucleases with different PAMs, plant systems, or base editors. Second, few cell types currently have both sufficient off-target sites and corresponding AlphaGenome tracks. Because chromatin accessibility is cell-type-specific, performance in cell types lacking matched data remains uncertain. Finally, sites undetected by GUIDE-seq were treated as negatives, though these are putative rather than confirmed negatives, as non-detection may reflect assay sensitivity or biological variability. If some of these undetected sites are in fact positives but are labeled and used as negatives, they may affect both model training and evaluation. During training, they introduce incorrect supervision and may hinder the model from learning patterns associated with true off-target activity. During evaluation, such sites are treated as negatives, so assigning them high scores increases the apparent false-positive burden and reduces measured precision, while they cannot contribute as true positives to recall. Consequently, AUC-PR may underestimate performance relative to the unknown true labels.

## Conflicts of interest

T.C. has a sponsored research collaboration with Cellectis and is an advisor to AaviGen, AstraZeneca, Novo Nordisk, and T-knife Therapeutics.

## Funding

Funded by the Deutsche Forschungsgemeinschaft (DFG, German Research Foundation) – Project-ID 499552394 – SFB 1597 to R.B. and T.C.

We appreciate the support by the Deutsche Forschungsgemeinschaft (DFG, German Research Foundation) through the research training group “BioInMe” 322977937/GRK2344 to D.R. and R.B.

This research was funded by the Deutsche Forschungsgemeinschaft (DFG, German Research Foundation) under grant number 539134284, through EFRE (FEIH_2698644) and the state of Baden-Württemberg.

We acknowledge support by the Open Access Publication Fund of the University of Freiburg.

## Data availability

All datasets and code required to reproduce the results in the manuscript are publicly available at: https://github.com/domonik/CRISCrossPaper

In addition, the CRISCross model is available separately as a Python package at: https://github.com/domonik/CRISCross (47)

## Author contributions statement

W.J.P. implemented the dataset processing pipeline, extended state-of-the-art models with late merging, designed and conducted experiments on tabular and sequence models, performed feature analyses, and wrote the manuscript. D.R. designed and implemented CRISCross and CRISPRAT, conducted the corresponding experiments, created plots, and wrote the manuscript. M.K. contributed to writing the manuscript. A.M. created a Windows GUI version of the model. M.U. revised the manuscript. T.B. revised parts of the manuscript. T.C. revised the manuscript, supervised the study, and secured funding. V.D.T. supervised the study and wrote the manuscript. R.B. supervised the study, wrote the manuscript, and secured funding.

All authors contributed to discussions and approved the final manuscript.

## Key Points

- Epigenetic features contribute to CRISPR-Cas9 off-target prediction, but their predictive value cannot be effectively exploited through simple feature integration.
- CRISCross introduces nucleotide-level cross-attention to explicitly model interactions between sequence and epigenetic context, enabling more effective use of chromatin information.
- We propose CRISPRAT, a synthetic pretraining strategy based on genome-derived sgRNA-off-target pairs, which improves generalization across guide RNAs, datasets, and cell types, enabling competitive cross-cell-type prediction performance.

## Declaration of AI-assisted technologies in the manuscript preparation process

During the preparation of this work, the authors used Chat-GPT (OpenAI) to improve the language and readability of the manuscript. After using this tool, the authors reviewed and edited the content as needed and take full responsibility for the content of the article.

## Notes

### Competing Interest Statement

Toni Cathomens has a sponsored research collaboration with Cellectis and is an advisor to AaviGen, AstraZeneca, Novo Nordisk, and T-knife Therapeutics.

