## Supplementary Information for "CRISCross: Integration of Epigenetic Information for Deep-Learning-Based Prediction of CRISPR-Cas9 Off-Targets Across Cell Types"

### Supplementary Results

#### No performance improvement is observed with increasing context size

To evaluate the effect of incorporating additional sequence and contextual information, we extended the input beyond the 23-nt region by including flanking sequences of 128 and 512 nucleotides. We compared these extended-context models with the 23-nt baseline across different feature sets, namely sequence-only, experimental epigenetic features, and AlphaGenome-predicted features.

Across all feature sets, increasing the context size did not result in statistically significant improvements compared to the corresponding 23-nt models (Fig. S6). For sequence-only models, performance showed a slight increase from an AUC-PR of 0.741 to 0.764 with larger context windows. In contrast, models incorporating experimental or AlphaGenome-derived features exhibited a gradual decline in performance as context size increased.

These observations are consistent with our previous analysis of epigenetic feature contributions, which revealed distinct scale-dependent patterns. In particular, AlphaGenome-predicted ATAC-seq signals exhibited their largest effect sizes and strongest statistical significance within a narrow window centered on the target site (23 nt), while histone modification signals exhibited maximal effect sizes at substantially larger genomic scales ($\sim$8 kb).

Despite the apparent informativeness of histone marks at larger scales, extending the input context provided to the model did not improve predictive performance. This suggests that CRISPR-Cas9 off-target activity is primarily governed by localized sequence and chromatin accessibility signals. Naively expanding the input window introduces redundant and noisy features, dilutes target-specific information, and complicates the identification of relevant positions.

While we cannot fully exclude the possibility that histone features may become beneficial at substantially larger context sizes, effectively leveraging such long-range information is non-trivial and would likely require specialized architectures, such as Caduceus (1), which are designed to process kilobase-scale genomic sequences efficiently within computational constraints.

#### Robustness to Negative Sampling Strategy

To assess whether our cross-cell-type results depend on the specific negative-sampling scheme used for fine-tuning, we repeated the T-cell dataset subsampling with five additional random seeds and separately constructed a hard-negative variant in which the mismatch distribution of sampled negatives was closer to that of the GUIDE-seq-detected positives (Fig. S8). Across the five runs using the additional random seeds, performance closely matched our original results, indicating that the Amodel is robust to this source of variability. With hard-negative sampling, overall performance on the K562 test set decreased for both feature sets. Nonetheless, the model incorporating AlphaGenome ATAC-seq features still outperformed the sequence-only model (0.483 vs. 0.441). The drop in overall performance likely reflects two related factors: a mismatch between the harder training negatives and the comparatively easier K562 test negatives, and an increased risk that hard negatives, being more similar to true positives, include undetected true off-target sites mislabeled as negatives due to assay sensitivity limits, introducing label noise into training.

### Supplementary Methods

#### Problem formulation

We formulate the CRISPR-Cas9 off-target prediction as a supervised binary classification problem. Given a single guide RNA (sgRNA) sequence, a candidate genomic sequence, and the corresponding epigenetic signal tracks, the objective is to predict whether the site is cleaved by the Cas9 nuclease.

Let $s$ denote the 23-nucleotide-length sgRNA sequence combined with the on-target PAM, and $t$ the candidate genomic sequence centered on the potential off-target site, which may include additional upstream and downstream context and therefore has variable length $L$. In addition, each candidate site is associated with position-resolved epigenetic signals $E$, consisting of multiple tracks aligned in the same genomic region as $t$. The prediction task is to learn a parameterized function: $f_\theta(s,t,E) \to y \in [0,1]$, where $\theta$ denotes the model parameters and $y$ represents the predicted probability that the candidate site is a true off-target. During training, ground truth labels are provided as $y \in \{0,1\}$, indicating experimentally validated cleavage events

### SHAP values

Shapley values are a way to explain the prediction of a model $f(x)$ for a specific input $x$. SHAP values are based on measuring the model prediction $v_x(S)$ made by model $f$ for data point $x$ when using only the features in the subset $S \subseteq M$ of the set of all features $M$. Here, $v_x(S)$ can be defined in different ways. In the strongest definition (see Lundberg and Lee 2017 (2)) it is defined by defining $f_S$ as the model trained using only the features in $S$ and setting $v_x(S) = f_S(x_S)$, where $x_S$ represents the values of the input features in $S$. Thus $v_x(S)$ is the prediction for data point $x$ made by a model trained using only the features in $S$. The need to retrain the model can, for example, be avoided by defining $v_x(S)$ as the expected model prediction when using the feature set $S$ and marginalizing over all other features $C = M \backslash S$. SHAP as proposed Lundberg and Lee (2) and used here, relies on different estimates. However, similar principles apply. We refer the reader to Lundberg and Lee (2) for full details.

A Shapley value $\phi_x(i)$ for a specific feature $i$ and data point $x$ is generated from assessing the performance gain or loss of a specific feature $i \in M$ when added to a subset $S$, i.e., the value $v_x(S \cup \{i\}) - v_x(S) = f_{S\cup\{i\}}(x_{S\cup\{i\}}) - f_S(x_S)$. Then all possible subsets $S \subseteq M\backslash\{i\}$ are considered for calculating $\phi_x(i)$, normalized such that every split size has the same contribution:

$$\begin{aligned}
\phi_x(i) &= \frac{1}{m} \sum_{0\le s\le m-1} \binom{m-1}{s}^{-1} \\
&\quad \cdot \sum_{\substack{S\subseteq M\backslash\{i\} \\ |S|=s}} \left[v_x(S\cup\{i\}) - v_x(S)\right] \\
&= \sum_{S\subseteq M\backslash\{i\}} \frac{1}{|M|} \binom{|M|-1}{|S|}^{-1} \left[v_x(S\cup\{i\}) - v_x(S)\right] \\
&= \sum_{S\subseteq M\backslash\{i\}} \frac{|S|!(|M|-1-|S|)!}{|M|!} \left[v_x(S\cup\{i\}) - v_x(S)\right].
\end{aligned}$$

where $m = |M|$ is the total number of features and $\binom{m-1}{s}$ is the number of possible subsets $S \subseteq M\backslash\{i\}$ that have the size $s = |S|$. The Shapley values $\phi_x(i)$ for each feature $i \in M$ satisfy the local accuracy property in that they are an additive explanation for the model prediction $f(x)$ on input $x$:

$$f(x) = v_x(\emptyset) + \sum_{i\in M} \phi_x(i)$$

Beeswarm plots are an efficient way to display SHAP values. Here, the features are ordered by mean absolute Shapley value (i.e., by $\frac{1}{|X|}\sum_x |\phi_x(i)|$). For each feature row, a dot represents one data point $x$ and is positioned along the x-axis according to $\phi_x(i)$. When several points overlap in the same area, they are distributed along the y-axis. The color of each dot represents the value $x_i$ of that feature indicating which feature values increase or decrease the model prediction score.

**Epigenetic feature processing** For each feature, we computed both per-nucleotide signal intensities and summary statistics over multiple genomic ranges. Per-nucleotide signals for experimental features were extracted using pyBigWig for each 23-nt off-target site and surrounding context regions of 128 and 512 nucleotides. Summary statistics (mean, median, minimum, and maximum) were computed for the 23 nt off-target site as well as symmetric flanking windows of sizes 128, 256, 512, 1024, 2048, 4096, and 8192 bp centered on the off-target site.

The selected epigenetic markers correspond to the epigenetic features for which experimental datasets were available and that could also be predicted by AlphaGenome for the T-cell context, ensuring a consistent feature space for direct comparison between experimental and predicted signals.

To support CRISPRAT pretraining, genome-wide predicted ATAC-seq tracks were precomputed. The genome was partitioned into chunks of $2^{16}$ nucleotides, and for each chunk, the AlphaGenome API was queried using a 1 Mbp window centered on the chunk for both T-cell and K562 ontology terms. The resulting predictions were subsequently stitched together to reconstruct genome-wide feature tracks.

### Model training and implementation details

**Optimization:** All models were trained using the AdamW optimizer with parameters $\beta_1 = 0.9$, $\beta_2 = 0.999$, and $\epsilon = 10^{-8}$. A learning rate of $10^{-4}$ was used. A linear warmup followed by cosine decay was applied to the learning rate schedule during pretraining. A dropout rate of 0.2 was used to mitigate overfitting.

**Fine-tuning procedure:** Models were fine-tuned for up to 100 epochs using a batch size of 256. Early stopping was applied based on validation AUC-PR with patience of 10 epochs. For each training split, 20% of the training data was held out as a validation set for model selection. To fully leverage the T-cell dataset in the cross-cell-type evaluation, CRISCross was trained for the average number of steps determined from prior experiments (Suppl. Fig. S9). Additionally, stochastic weight averaging was applied from step 60 onward, as validation performance plateaued around this point.

**Loss function and class imbalance:** CRISPR off-target prediction datasets are highly imbalanced due to the large number of potential off-target sites used as negatives and the relatively small number of experimentally validated cleavage sites used as positives. In our datasets, the imbalance ratio is approximately 1:40 for the T-cell dataset and 1:155 for the K562 dataset.

The model was trained using binary cross-entropy (BCE) loss. To counteract class imbalance, a weighted random sampler was used during training only to construct mini-batches with approximately equal proportions of positive and negative samples. The sampling weights were assigned based on the class frequencies in the training set, giving higher sampling probabilities to samples from the underrepresented positive class. This resulted in a roughly balanced (1:1) class distribution within the training mini-batches, while the original class distribution is retained for validation and test data.

**Pretraining setup:** Both CRISPRAT and dataset-specific pretraining were performed using the same optimizer and learning rate. Pretraining was conducted with a base batch size of 2048 and gradient accumulation to achieve an effective batch size of 25,600. CRISPRAT pretraining was performed for 15,000 steps, while dataset-specific pretraining was conducted for 50,000 steps. The learning rate schedule used linear warmup followed by cosine decay, with 2,000 warmup steps for CRISPRAT and 5,000 warmup steps for dataset-specific pretraining.

For CRISPRAT, the training objective combines binary cross-entropy (BCE) loss for masked token prediction with a mean squared error (MSE) regression loss for thermodynamic prediction. In contrast, dataset-specific pretraining uses only the BCE loss.

**Model configuration:** CRISCross was configured with an embedding dimension of 512, used consistently across token embeddings, self-attention, and cross-attention layers. Both the self-attention and cross-attention sublayers use 4 attention heads and a feed-forward MLP with hidden dimension 4 times the embedding dimension (2048), with a dropout rate of 0.2 applied throughout. Sequences were tokenized using overlapping 3-mers. Three interaction blocks were used by default, each consisting of independent self-attention encoding of the sgRNA and genomic candidate representations, followed by bidirectional cross-attention restricted to the central 23-nt genomic window. As an ablation, the cross-attention join was compared against an alternative fusion strategy for combining the sgRNA and genomic candidate representations at each block: element-wise addition. Unless otherwise stated, cross-attention was used and this configuration was applied in all experiments.

### Supplementary Figures

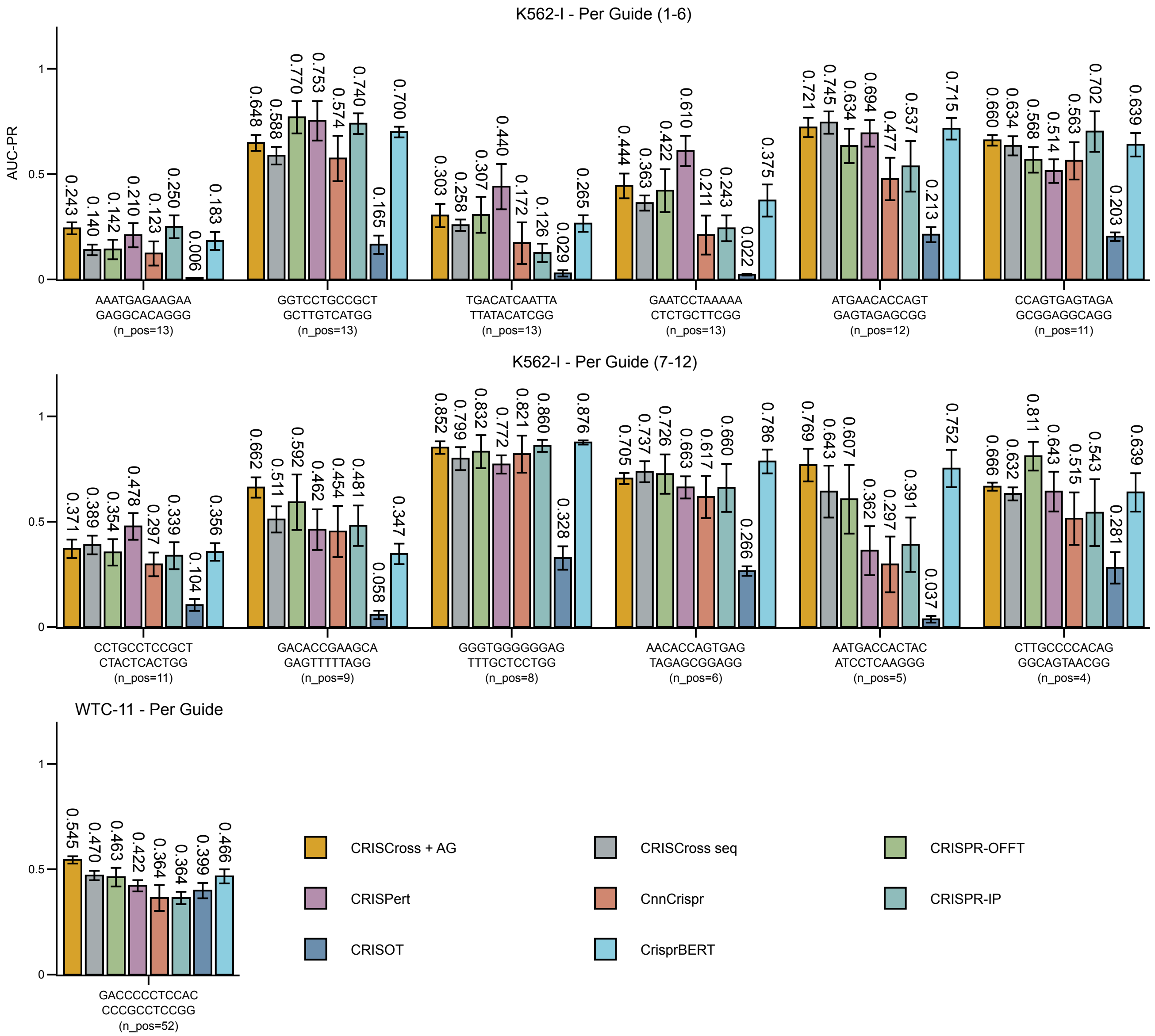


**Fig. S1.** Mean per-sgRNA AUC-PR and across seed standard deviation on K562-I (rows 1 and 2) and WTC-11 (row 3), with guides ordered by the number of true positive off-target sites (n_pos.)

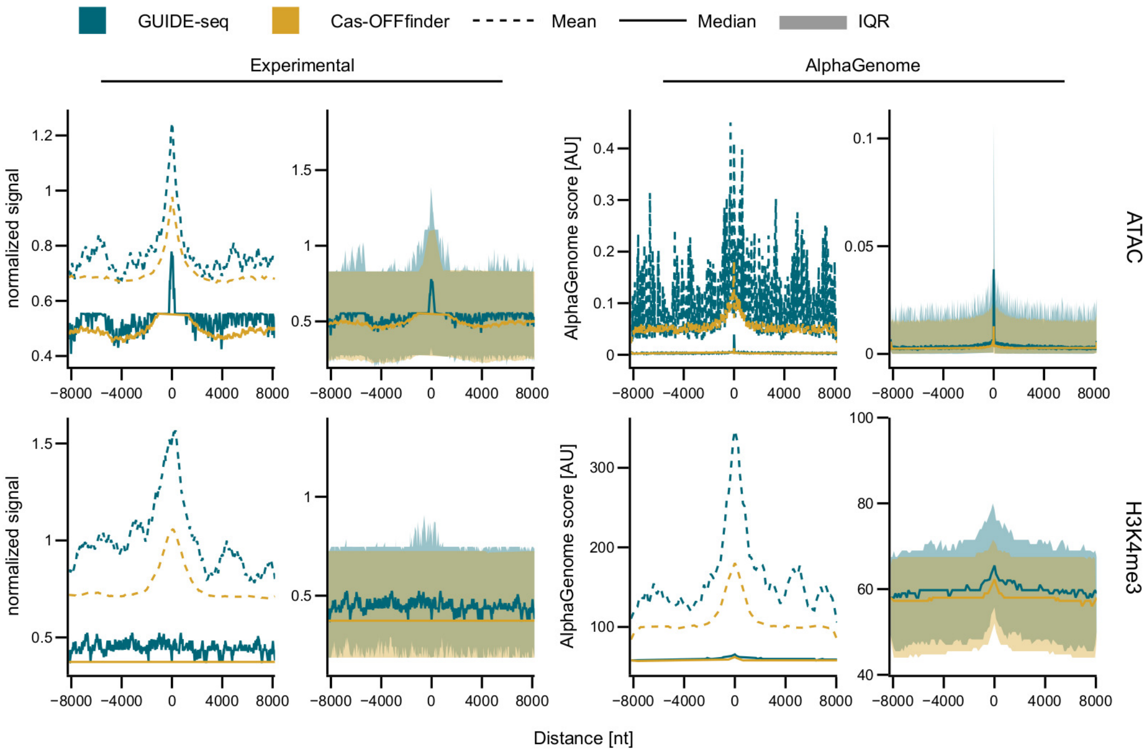


**Fig. S2.** Epigenetic signal profiles around off-target sites. Shown are aggregated signal distributions centered on off-target loci (0 nt) for experimental datasets (left) and AlphaGenome predictions (right), across ATAC and H3K4me3 marks. GUIDE-seq sites are shown in teal and Cas-OFFinder sites in orange. Dashed lines indicate the mean, solid lines the median, and shaded regions the interquartile range (IQR). While mean profiles suggest differences between site classes, broad IQRs and deviations between mean and median highlight high variability and skewness, reducing overall separability between groups.

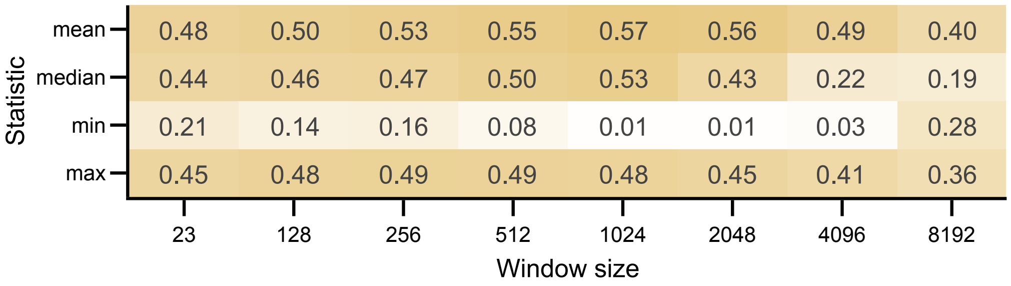


**Fig. S3.** Pearson correlation of the summary statistics for the AlphaGenome predicted and experimentally detected ATAC seq tracks.

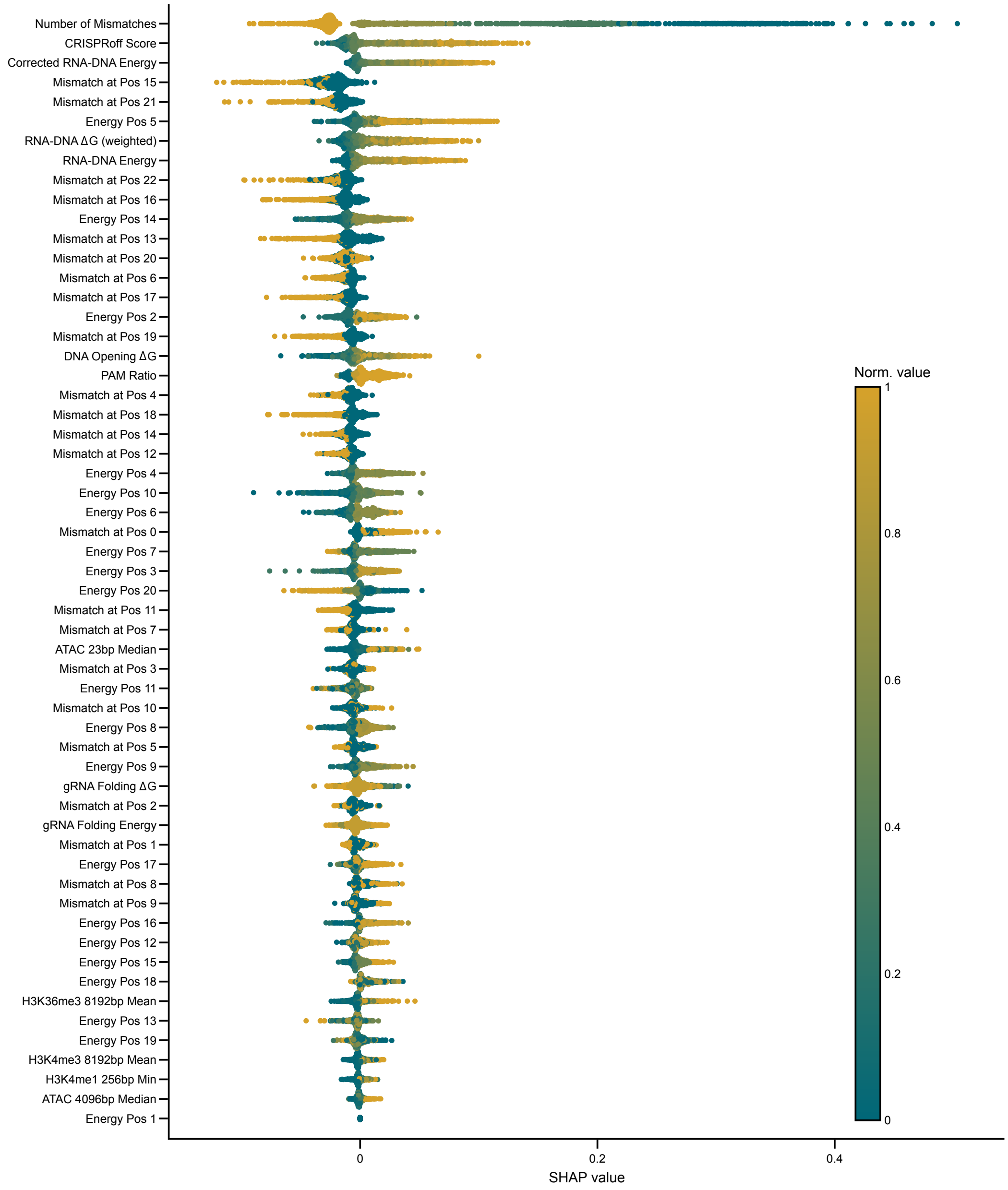


**Fig. S4.** Full set of SHAP values from the TabPFN evaluation run.

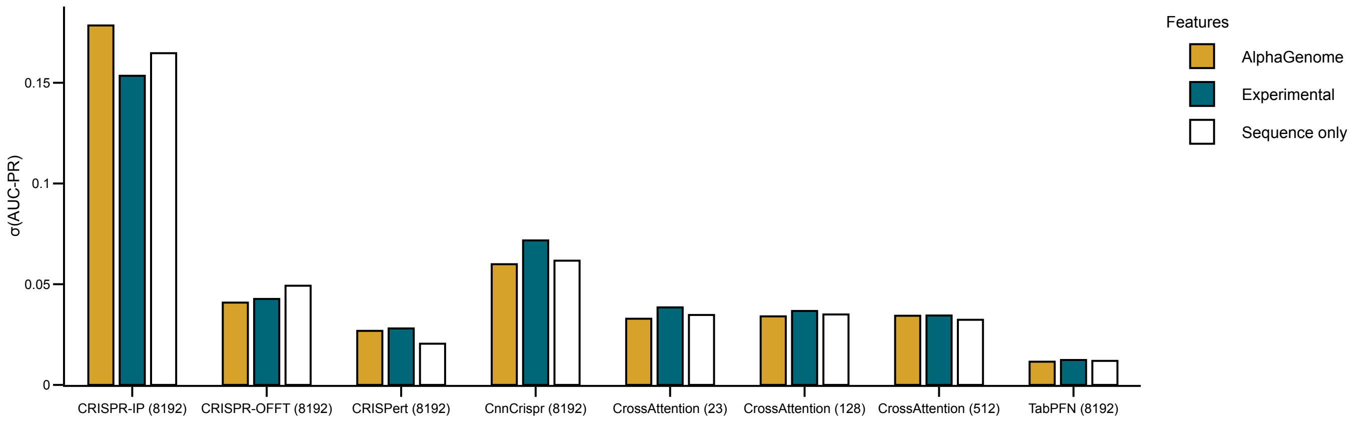


**Fig. S5.** Standard deviation of AUC-PR values across different seeds in the leave-one-guide-out setting for the T-cell dataset. Numbers in brackets show the maximum context window size of used features.

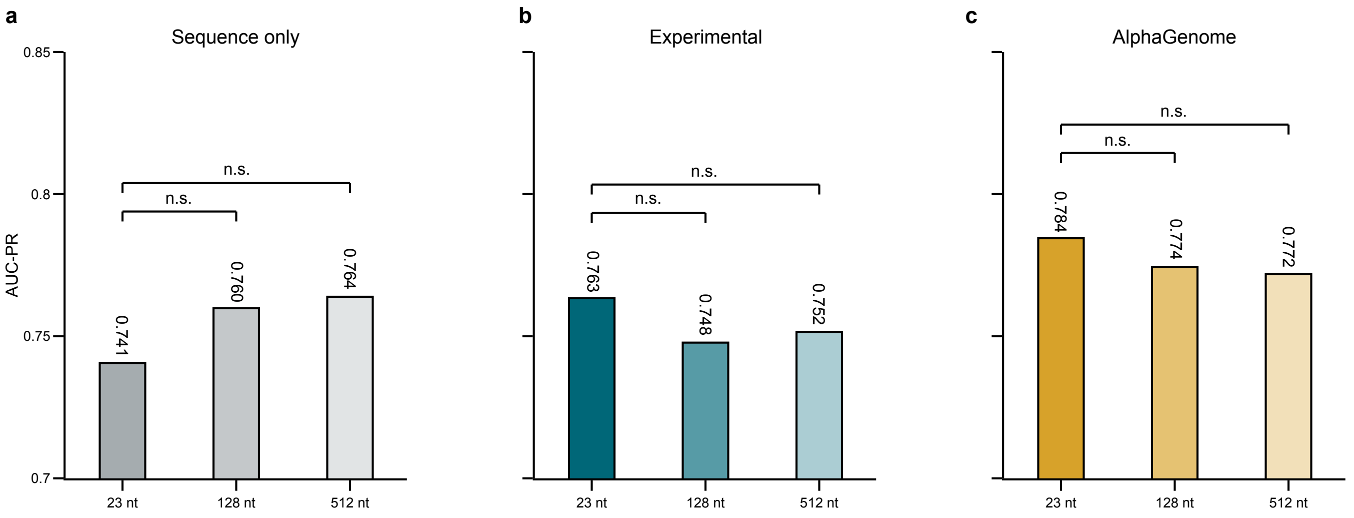


**Fig. S6.** Effect of context size on CRISCross performance across feature sets. AUC-PR is shown for CRISCross models trained with sequence-only (a), experimental epigenomic (b), or AlphaGenome-derived (c) input features and evaluated at context sizes of 23, 128, and 512 nt. Within each feature set, the 128-nt and 512-nt models are compared against their corresponding 23-nt baseline using a Wilcoxon signed-rank test with Holm-Bonferroni correction. Bar values indicate mean AUC-PR across cross-validation splits. Significance levels: * $p < 0.05$, ** $p < 0.01$, *** $p < 0.001$, n.s. = not significant.

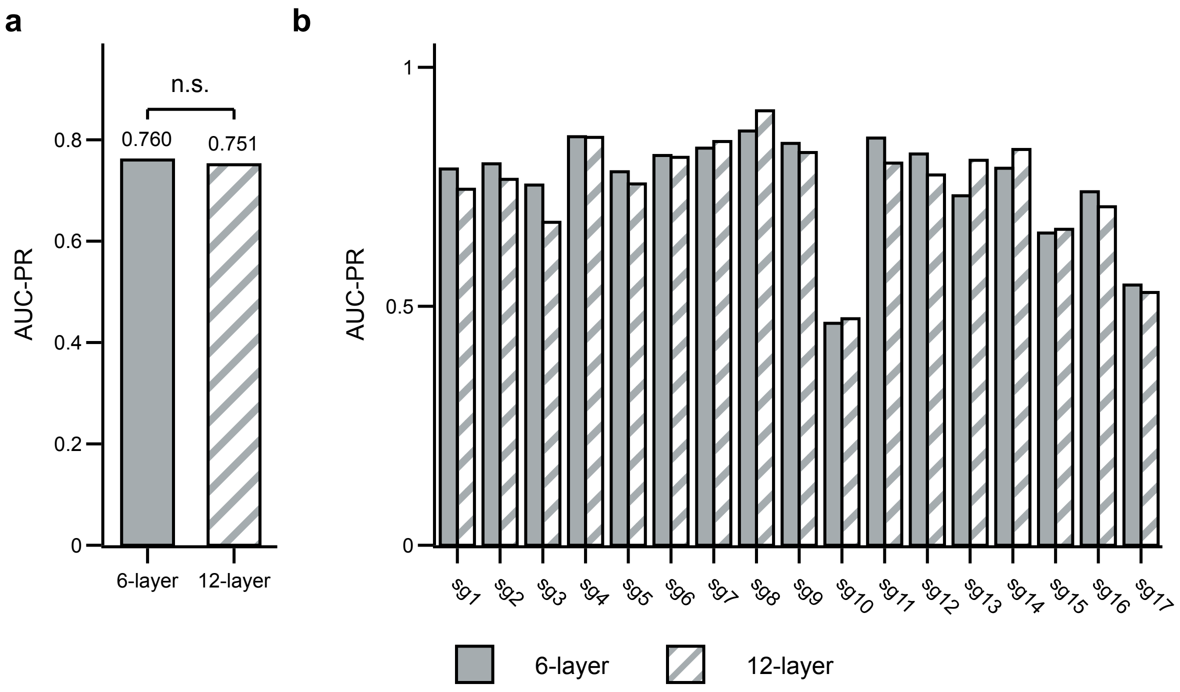


**Fig. S7.** Reducing CRISPert from 12 to 6 transformer layers does not significantly affect predictive performance. (a) Overall AUC-PR for the 6-layer (gray) and 12-layer (patterned) CRISPert models, averaged across all test sgRNAs. Mean AUC-PR was nearly identical between architectures (0.760 vs. 0.751), and the difference was not statistically significant (n.s.). (b) Per-guide breakdown of AUC-PR for both model depths across each of the 17 individual test sgRNAs (sg1–sg17), showing that the 6-layer and 12-layer models track each other closely at the level of individual guides as well, indicating that the additional model depth provides no consistent benefit.

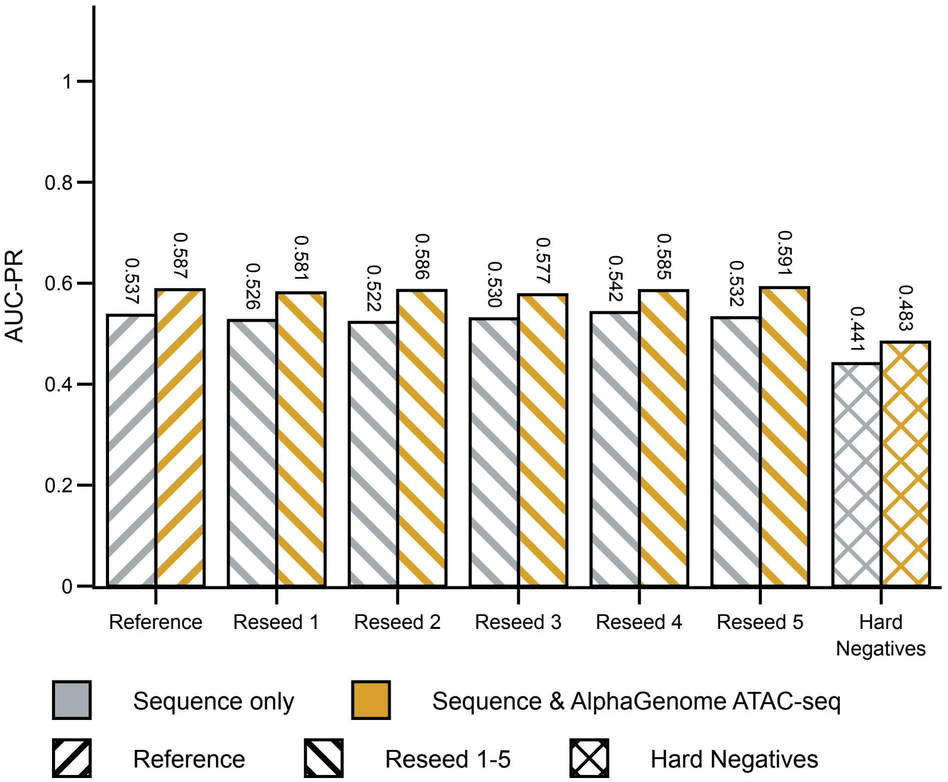


**Fig. S8.** Robustness of CRISCross to fine-tuning negative-sampling choices. AUC-PR on the K562-I test set for CRISCross fine-tuned on the original T-cell dataset, on five independently re-sampled T-cell datasets (different random seeds for negative sampling), and on a hard-negative T-cell dataset in which negative mismatch distributions were matched to those of GUIDE-seq-detected positives, shown for both sequence-only and AlphaGenome ATAC-seq-augmented models. Performance is consistent across the re-sampled seeds, indicating robustness to the specific negative sample drawn during fine-tuning. Performance decreases for both feature sets under hard-negative sampling, but the AlphaGenome ATAC-seq model continues to outperform the sequence-only model (0.483 vs. 0.441).

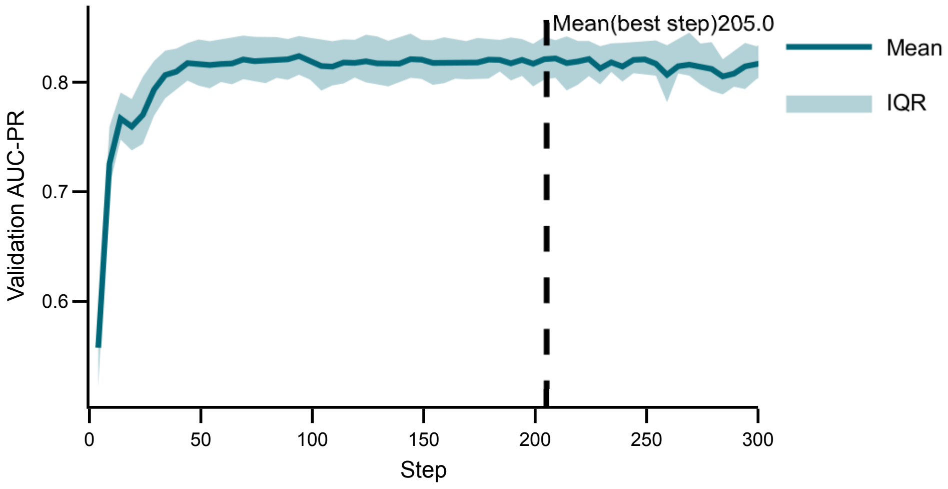


**Fig. S9.** Validation performance of CRISPRAT models in the leave-one-guide-out setting. Mean and interquartile range of AUC–PR values are plotted against training steps, with the dashed line indicating the mean step at which the best validation AUC–PR was achieved across all runs.
